# Charting the emergent organotypic landscape of the mammalian gut endoderm at single-cell resolution

**DOI:** 10.1101/471078

**Authors:** Sonja Nowotschin, Manu Setty, Ying-Yi Kuo, Vincent Lui, Vidur Garg, Roshan Sharma, Claire S. Simon, Nestor Saiz, Rui Gardner, Stéphane C. Boutet, Deanna M. Church, Pamela A. Hoodless, Anna-Katerina Hadjantonakis, Dana Pe’er

## Abstract

To comprehensively delineate the ontogeny of an organ system, we generated 112,217 singlecell transcriptomes representing all endoderm populations within the mouse embryo until midgestation. We employed graph-based approaches to model differentiating cells for spatio-temporal characterization of developmental trajectories. Our analysis reveals the detailed architecture of the emergence of the first (primitive or extra-embryonic) endodermal population and pluripotent epiblast. We uncover an unappreciated relationship between descendants of these lineages, before the onset of gastrulation, suggesting that mixing of extra-embryonic and embryonic endoderm cells occurs more than once during mammalian development. We map the trajectories of endoderm cells as they acquire embryonic versus extra-embryonic fates, and their spatial convergence within the gut endoderm; revealing them to be globally similar but retaining aspects of their lineage history. We observe the regionalized localization of cells along the forming gut tube, reflecting their extra-embryonic or embryonic origin, and their coordinate patterning into organ-specific territories along the anterior-posterior axis.

## Introduction

How individual cells make fate decisions within populations, and how populations of cells are reproducibly patterned to coordinately build tissues in a stereotypical fashion are longstanding questions in biology. A notable feature of mammalian embryos is their association with extra-embryonic tissues, which provide the interface with the mother, as well as nutritive support ^1^. The earliest cell fate decisions taking place in mammals serve to segregate embryonic (epiblast) from extra-embryonic (trophoblast, and primitive endoderm) lineages. During mammalian development, endoderm emerges at two distinct times (**Fig. 1**). Cells with a primitive (also known as extra-embryonic) endoderm (PrE) identity arise in the blastocyst stage embryo at mouse embryonic day (E)3.5 ^2,3^. Around E7.0, the definitive endoderm (DE) is specified from epiblast (EPI) at gastrulation ^4^. The gut endoderm, the precursor of the respiratory and digestive tracts, and their associated organs such as the thymus, thyroid, liver and pancreas, is one of the first tissues to be established in bilaterian embryos ^1,5,6^. Our previous studies revealed that the gut endoderm of the mouse embryo arises from cells of two different origins; comprising descendants of PrE and DE (**Fig. 1A**) ^7–9^. However, despite their distinct origins and locations, embryonic (gut endoderm) and extra-embryonic (visceral, parietal and yolk sac) endoderm share many molecular features hampering marker-based discrimination ^10,11^.

**Figure 1:**
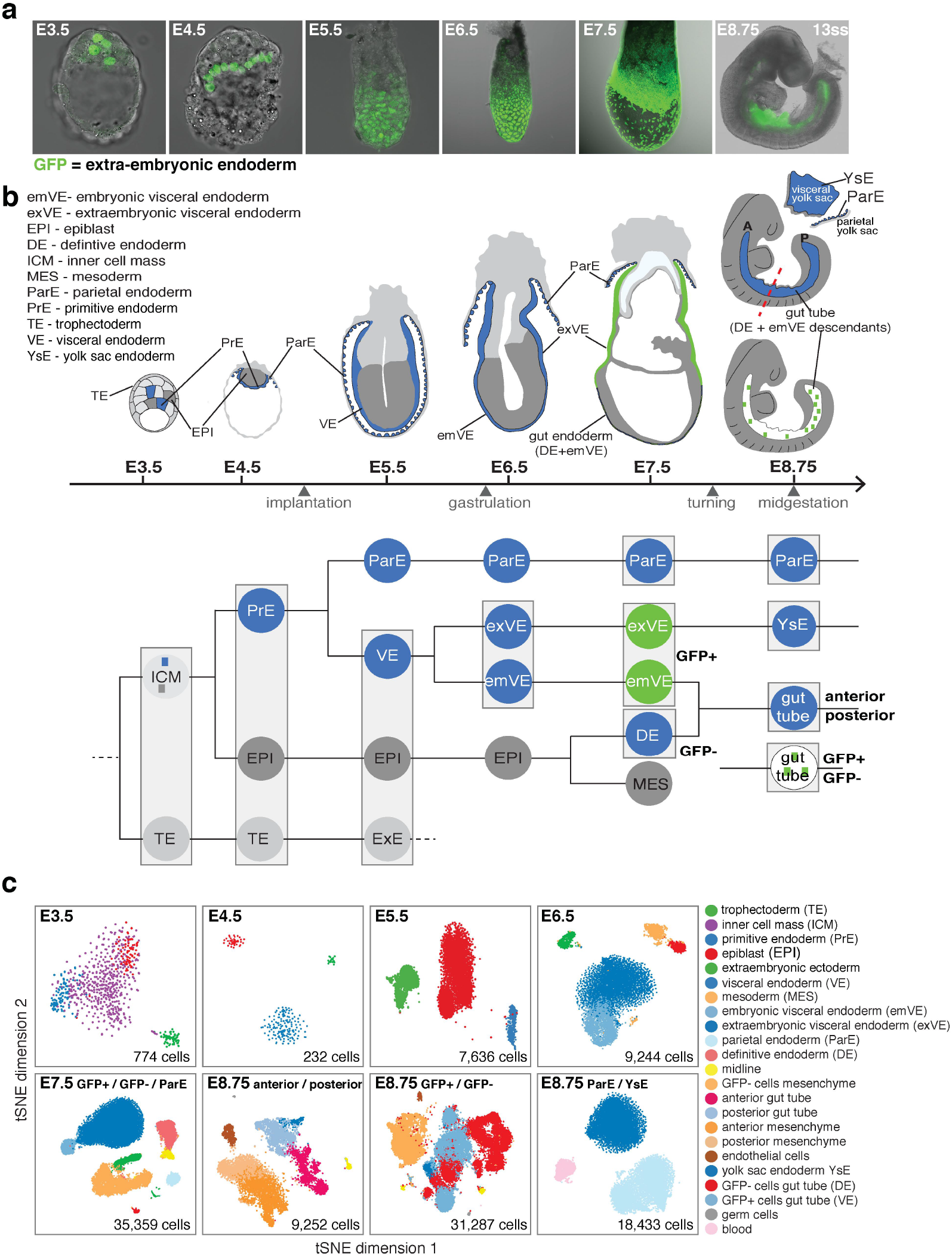
Single cell map of the endoderm from blastocyst to midgestational in the mouse. **a,** Distribution of extra-embryonic endoderm cells (GFP/green) from the blastocyst (E3.5) to midgestation (E8.75, 13ss) demarcated using Pdgfra^H2B-GFP^ (pre-implantation stages) and Afp-GFP (post-implantation stages) reporters. Extra-embryonic endoderm (PrE and VE derivatives) cells contribute to the gut tube of the E8.75 embryo. **b,** Schematic of cell lineage tree, experimental design and the 13 single-cell libraries collected across sequential stages (grey boxes) with all sampled in duplicate or triplicate. **c,** tSNE plots of collected libraries for each time-point with each dot representing a single cell. Phenograph was used to identify clusters of cells, color-coded by cell type annotation based on expression of known markers.

To map the landscape of the mammalian gut endoderm, we sought to systematically and comprehensively characterize the transcriptional profiles of all endoderm cell populations in the mouse embryo; from the blastocyst, when the first endodermal cells appear, to midgestation, when the gut endoderm comprises an internal tube, which becomes regionally patterned as its cells acquire distinct organ identities along its anterior-posterior (AP) axis. For this, we generated 112,217 single-cell transcriptomes using the 10x Chromium single-cell 3’ RNA-seq platform. Cells were collected from wild-type embryos for all stages analyzed, complemented at later stages (E7.5 and E8.75) with an extra-embryonic endoderm-specific reporter for the isolation of PrE and DE descendants (**Fig. 1B and Extended Data Fig. 1**). To characterize developmental trajectories spanning discrete time-points, we developed Harmony, a graph-based framework that seamlessly connects consecutive time-points. We use Harmony in conjunction with our differentiation trajectory inference algorithm, Palantir ^12^ to infer developmental trajectories, lineage relationships and the emergence of a stereotypical patterned organization within the gut tube. Pseudo-spatial mapping was performed using diffusion maps ^13^, allowing us to map cell positions without *a priori* information of marker localization. These positional assignments were validated by bulk RNA-seq data of regionally micro-dissected tissues.

Our analyses reveal the detailed architecture of key mammalian lineage specification events, notably the emergence of PrE versus pluripotent epiblast from an inner cell mass progenitor pool within the blastocyst. We uncover an unappreciated lineage relationship between the epiblast and visceral endoderm in the early (E5.5) post-implantation embryo, before the onset of gastrulation, which we validate using lineage-tracing in embryos. Furthermore, we have compiled a detailed map of the trajectories adopted by endoderm cells as they acquire embryonic versus extra-embryonic fates along the proximal-distal axis of the early embryo, identifying key bifurcation and convergence points of embryonic and extra-embryonic tissues leading to the establishment of distinct territories along the AP axis of the E8.75 gut tube presaging the emergence of the endodermal organs. We observe, and validate in embryos, the stereotypical regionalized localization of cells along the gut tube reflecting their origin, and subsequent patterning into organ-specific territories along the AP axis. Despite being globally similar, our analyses reveal that gut endoderm cells retain aspects of their lineage history and can be distinguished by a core set of genes. In sum, this study, in a mammalian model, provides the first comprehensive transcriptional characterization of the ontogeny of an organ system, the gut endoderm.

## Results

To achieve a comprehensive isolation and characterization of all endodermal cell types between the blastocyst and midgestation (**Fig. 1a-b**), we isolated cells from sequentially-staged wild-type mouse embryos between E3.5 and E8.75, including associated endoderm-containing extra-embryonic tissues. Cohorts of cells representing each population sampled were processed for scRNA-seq library preparation using the 10x Genomics Chromium platform (**Fig. 1b,** libraries demarcated as grey boxes). Due to their small size, low total cell number and difficulties isolating tissue-layers at pre-implantation (E3.5-E4.5) through early post-implantation (E5.5), whole embryos were used for single-cell isolations. Embryos from later stages (E6.5-E8.75) are larger, comprise more cells, and thus endodermal tissue layers were isolated for cell-type enrichment, (endoderm comprises 3.5% of E8.75 embryos, **Extended Data Fig. 1a**).

Since the E8.75 gut tube comprises cells of two distinct origins ^7,9^, we also incorporated a lineage-tracing experiment into our study. Using an *Afp-GFP* reporter ^14^ to identify cells having an extra-embryonic (PrE/VE) origin and distinguish them from definitive endoderm (DE) neighbors having an embryonic (epiblast, EPI) origin, we isolated GFP-positive (extra-embryonic) and GFP-negative (embryonic) populations by flow cytometry after tissue dissociation at E7.5 and E8.75 (**Fig. 1b, Extended Data Fig. 1b, Supplementary Fig. 1**). We profiled 13 samples, each collected in duplicate or triplicate, representing a total of 112,217 cells (**Fig. 1c, Supplementary Fig. 2**). We processed each sample using our processing pipeline ^12,15^ (**Extended Data Fig. 2a**), verified replicates for each time-point were comparable (**Supplementary Fig. 2**), and then combined all samples for each time-point. To identify clusters at each time-point, cells were clustered using Phenograph ^16^, with identities assigned based on marker gene expression and visualized using tSNE ^17,18^. We are able to recover all the known cell types across the different stages of developing endoderm thus providing a comprehensive characterization of mouse endoderm development (**Fig. 1c**).

To demonstrate that scRNA-seq data was representative, and that dissociation did not alter the transcriptome of cells, we generated bulk RNA-seq data of dissociated cells or whole tissue from isolated E8.75 gut tubes (**Extended Data Fig. 1b**). Gene expression counts from both bulk RNA-seq samples highly correlated with one another, and with aggregated scRNA-seq expression counts (**Extended Data Fig. 2b**), indicating that our tissue isolation and dissociation procedure did not alter cell type proportions or transcriptional profiles.

Following recent successes in reconstructing developmental trajectories from scRNAseq ^19–26^, we aimed to organize cells along developmental trajectories that would elucidate when and how cell-fate decisions occur. However, connecting cells across distinct time-points presents challenges: (a) The extensive cell divisions associated with the developing embryo means we might fail to capture key intermediate stages between consecutive time-points, and (b) True biological differences between time-points are often confounded by batch-effects. Current batch-effect correction approaches ^22,27^ force similarity of cell states across batches, and hence can dilute important changes between time-points (**Extended Data Fig. 3f**). We therefore developed Harmony, a robust framework that connects cells across time-points by augmenting the nearest-neighbor graph (kNN graph), a construct encoding cell-cell similarities that underlies many scRNAseq algorithms ^16,27–29^. Harmony takes advantage of asynchrony present at each time-point, i.e., a subset of more mature cells in one time-point are relatively closer to another subset of least mature cells in the subsequent time-point, resulting in mutually-similar cells across time-points. Harmony harnesses these mutual nearest neighbors to construct the augmented kNN-graph that connects the time-points together (**Extended Data Fig. 3**). The augmented kNN-graph unifies the data so that it can then be provided as input to any kNN-graph based algorithm (**Extended Data Fig. 4a**) and data imputation using MAGIC ^30^.

To construct developmental trajectories we combined Harmony with Palantir^12^, our recent pseudo-time algorithm for modeling cell fate decisions. Rather than treating lineage decisions as discrete bifurcations, Palantir models cell fate choices as a continuous process: for each cell state, Palantir infers its probability to reach each of the terminal fates in the system. The entropy of these fate probabilities represents the differentiation potential – a measure of the degree of uncertainty in cell fate associated with a cell in a particular state. Regions along pseudo-time where changes in differentiation potential occur represent regions of lineage specification and commitment (See supplementary note on Palantir). The Palantir framework also enables a high resolution characterization of gene expression dynamics during lineage specification and comparison of these dynamics across lineages^12^. Thus, Palantir provides insight into when fate decisions occur, and what gene expression changes coincide with this timing.

### Detailed structure of a seemingly simple lineage decision: emergence of first endodermal population, the primitive endoderm

The pre-implantation period of mammalian development culminates in the formation of the blastocyst comprising three cell lineages^2^; trophectoderm (TE), which gives rise to the fetal portion of the placenta; epiblast (EPI), which gives rise to most somatic lineages, germ cells and some extra-embryonic mesoderm; primitive endoderm (PrE), which gives rise to the endodermal component of the visceral and parietal yolk sacs, and gut endoderm. Force directed layout following Harmony between E3.5 and E4.5 datasets illustrates the relationship between the blastocyst lineages (**Fig. 2a-b**). To highlight the dominant components of change, we projected the data onto diffusion components ^13,20,23,31,32^, and measured the average distance of the three lineages from the bipotent inner cell mass (ICM) (**Fig. 2b**). TE cells were substantially farther away (13.9) from ICM compared to either EPI (0.41) or PrE (2.0). This suggests that TE-vs.-ICM, the first cell fate decision, is complete by E3.5, prior to ICM cells making a PrE-vs.-EPI choice, and that EPI cells are phenotypically closer to ICM cells, compared to PrE.

**Figure 2:**
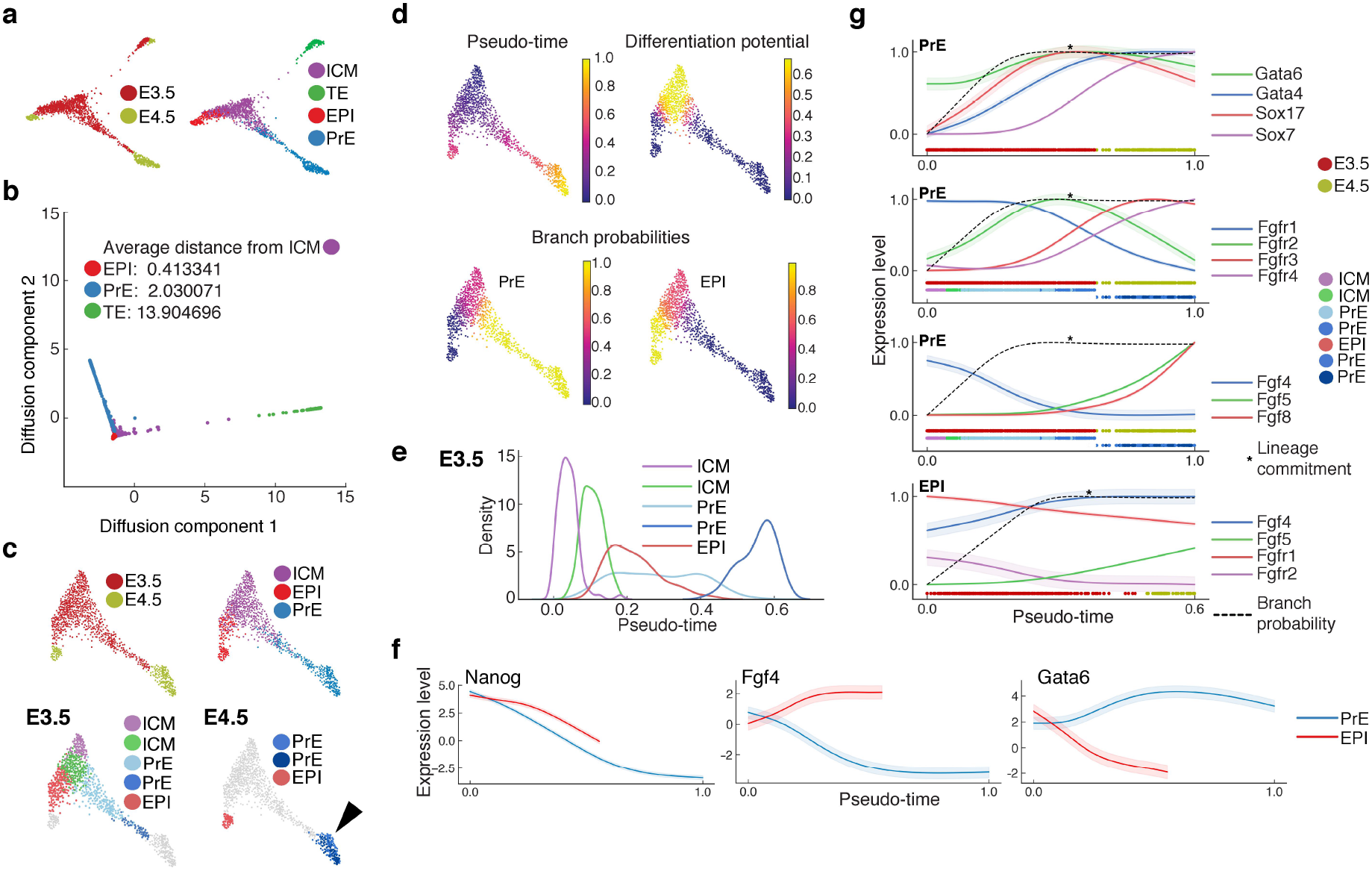
Lineage decisions in the pre-implantation mammalian blastocyst. The results shown were generated by pooling cells from the two replicates each for E3.5 and E4.5 and using Harmony augmentation. **a,** Force directed layout of E3.5 and E4.5 cells showing the relationships between the three blastocyst lineages. Cells are colored by Phenograph clusters. **b,** Plot showing the projection of E3.5 and E4.5 cells along the first two diffusion components. Distances between lineages were computed using multi-scale distances. **c,** Force directed layout of E3.5 and E4.5, following the removal of TE cells, showing the relationships between ICM, EPI and PrE lineages. In the different panels, cells are colored by time-point or Phenograph clusters. **d,** Palantir determined pseudo-time ordering, differentiation potential and branch probabilities of PrE and EPI cell lineages at E3.5 and E4.5. See supplement note on Palantir for more information. **e,** Distribution of E3.5 lineage cells along pseudo-time, each distribution represents cells from one Phenograph cluster. **f,** Gene expression trends along pseudo-time, computed using Palantir, for Nanog, Fgf4 and Gata6 during EPI and PrE lineage specification. **g,** Plots comparing gene expression trends along pseudo-time: Gata and Sox transcription factors, Fgf-receptors or -ligands along PrE or EPI lineage specification. Colors at the bottom of each panel represent the time-point and where applicable E3.5 and E4.5 Phenograph clusters. Dotted lines represent branch probabilities in commitment towards the respective lineages.

TE cells were excluded for subsequent analyses investigating the architecture of the lineage decision in which PrE and EPI arise (**Fig. 2c**). To pinpoint the decision-making region and characterize gene expression dynamics during commitment, we applied Palantir ^12^ using a Nanog^hi^ cell (ICM, still uncommitted) as the start (**Fig. 2d**). The change in differentiation potential and branch probabilities of cells suggests ICM lineage divergence into EPI and PrE occurs at E3.5, consistent with previous data ^33,34^ (**Fig. 2d, Extended Data Fig. 5a**). Consistent with this prediction, our analysis identified two ICM clusters; one likely representing uncommitted cells (purple) having equal propensity for PrE or EPI fates, and a second (green), which albeit still uncommitted, had started to specify towards PrE or EPI (**Fig. 2e, Extended. Data Fig. 5b, Supp. Table 1**). In addition, we also identified two PrE clusters at E3.5, likely representing nascent (light blue) and more mature (dark blue) populations, and one EPI cluster (red) (**Fig. 2c**), likely reflecting successive lineage maturation^35,36^. In contrast, by E4.5, we observed two distinct populations representing the EPI and PrE lineages, respectively (**Fig. 2c, Supp. Table 2**). Within the emergent PrE population, we observed two clusters (**Fig. 2c,** light and dark blue, black arrowhead), which, based on marker expression, represented nascent visceral (VE, orange arrowhead) and parietal (ParE, black arrowhead) endoderm derivatives of the PrE (**Fig. 2c, Extended Data Fig. 5c**).

ICM cell fate specification is driven in part by the lineage-specific transcription factors Gata6 and Nanog, which are co-expressed in the uncommitted ICM and required for, PrE and EPI, respectively^37–42^. While active across the ICM population^43–45^, FGF signaling, via FGF4^46–48^, is critical for PrE specification and exit from the naïve state or pluripotency (EPI). While the general localization of key transcription factors and signaling pathway components is known, the dynamics of their coordinated expression patterns at single-cell resolution across the maturing ICM population are unclear. We used Palantir to characterize expression trends for Nanog, Fgf4, and Gata6 along pseudo-time, as PrE and EPI identities emerged (**Fig. 2f**). The ratio of Gata6/Nanog closely tracked with EPI specification (as determined by Palantir’s branch probability) and is a strong descriptor of ICM fate specification, while the Gata6/Fgf4 ratio also tracked with EPI specification but trailed that of Gata6/Nanog (**Extended Data Fig. 5f**). These data also facilitated the precise ordering of markers during lineage-specification. In the PrE, for example, Gata6 is expressed in cells prior to commitment and maintained thereafter, whereas Sox17, Gata4 and Sox7 are sequentially activated ensuing commitment (**Fig. 2g, top panel**)^36^.

Mouse mutants in both *Fgfr1* and *Fgfr2* phenocopy embryos lacking *Fgf4*, exhibiting defects in PrE specification and exit from naïve pluripotency in the EPI^44–47^. *Fgfr1* single mutants exhibit a reduction in PrE cells, while *Fgfr2* mutants have no discernable lineage specification defect, supporting Fgfr1 as the main receptor driving ICM lineage specification. The coordinated or distinct actions of these receptors within emergent lineages has remained elusive. Palantir revealed that Fgfr1 is expressed throughout the early ICM (**Fig. 2g, Extended Data Fig. 5d and 5e**). Upon PrE specification, Fgfr1 was downregulated, accompanied by a transient activation of Fgfr2. Fgrf1 and Fgrf2 are coordinately expressed during PrE specification suggesting that although Fgfr1 is the main receptor, Fgfr2 might be required to augment its function (**Fig. 2g, second panel**). At E4.5, both Fgfr3 and Fgfr4 were expressed in PrE, but Fgfr1 and Fgfr2 are downregulated, suggesting that an FGF signal may be reiteratively used, but differentially transduced within a single lineage. Activation of a second phase of FGF signaling within the PrE, could be executed by Fgf5/Fgf8/Fgfr4 in the VE (**Extended Data Fig. 5e,** orange arrowheads), and Fgf3/Fgfr3 in ParE (**Extended Data Fig. 5e,** black arrowheads), respectively, with Fgf4/Fgfr1 being the main ligand/receptor combination within the EPI driving pluripotent state transitions (**Fig. 2g, Extended Data Fig. 5d and 5e,** green arrowheads). In addition, we identified a number of genes that correlate with the expression dynamics of Fgf4 in EPI, and Gata6/Gata4 in PrE along pseudo-time from E3.5 through E4.5 (**Extended Data Fig. 6a**). Immunofluorescent staining of the transcription factor Tcf7l1, one of the genes correlating with Fgf4 in the EPI, revealed EPI-specific expression at E4.5 (**Extended Data Fig. 6b**).

### Precocious differentiation of epiblast into visceral endoderm, before the onset of gastrulation

While EPI and PrE populations were represented as distinct populations at E4.5, by E5.5, we observed cells bridging the EPI and visceral endoderm (VE, descendant of PrE) populations (**Fig. 3a, Extended Data Fig. 7a,** black arrowhead). By contrast, no connection was observed between EPI (or VE) and extra-embryonic ectoderm (ExE, descendant of TE, **Extended Data Fig. 7a-b**). As in the blastocyst (**Fig. 2a-b**), ExE cells were phenotypically more distinct from EPI and VE at E5.5, based on projections of cells along the diffusion components and average pairwise distances between populations (**Extended Data Fig. 7a-b**). Furthermore, shortest paths between EPI and VE lineage cells indicated a presence of a continuum of cell states bridging EPI and VE identities (**Extended Data Fig. 7c**).

**Figure 3:**
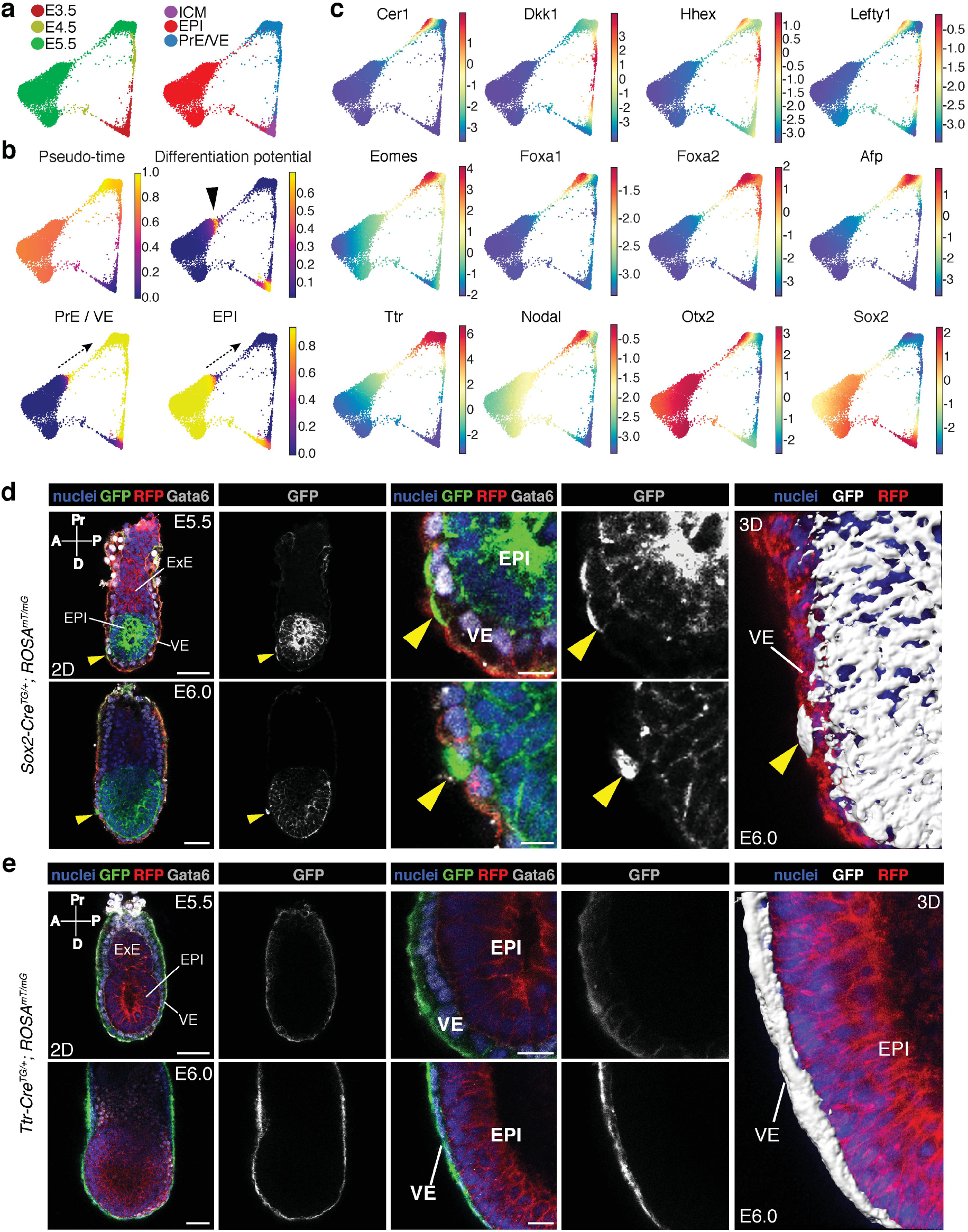
Trans-migration and trans-differentiation of epiblast into visceral endoderm at early post-implantation stages. The results shown were generated by pooling cells from all the replicates of E3.5, E4.5 and E5.5 and following Harmony augmentation. **a,** Force directed layouts of E3.5, E4.5 and E5.5 data depicting the relationships between the EPI and PrE/VE lineages. Harmony generated augmented affinity matrix to connect consecutive time-points was used to generate the visualization. Cells are colored by time points (left) and cell type labels (right). **b,** Plots showing Palantir pseudo-time, differentiation potential and branch probabilities of EPI and PrE/VE cell lineages at E3.5-E5.5. Black arrowhead and dotted arrows denotes EPI cells with high differentiation potential identified by Palantir representing a trans-differentiation of EPI to VE. **c,** Gene expression plots of AVE (Cer1, Dkk1, Hhex, Lefty1), VE (Eomes, Foxa1, Foxa2, Afp, Ttr), VE and EPI (Nodal, Otx2) and EPI (Sox2) markers along EPI and PrE/VE lineages from E3.5-E5.5. Cells colored based on marker expression of indicated gene. **d-e,** Laser confocal images of E5.5 and E6.0 Sox2-Cre^TG/+^; ROSA26^mT/mG^ (d) and Ttr-Cre^TG/+^; ROSA26^mT/mG^ (e) embryos immunostained for GFP, RFP (red fluorescent protein, membrane-localized tdTomato) and Gata6, a VE identity marker. 3D surface renderings of mGFP-expressing cells in E6.0 Sox2-Cre^TG/+^; ROSA26^mT/mG^ and Ttr-Cre^TG/+^; ROSA26^mT/mG^ embryos (panels on the right). Cell nuclei are stained with Hoechst and membranes are labeled with RFP. Yellow arrowheads point to epiblast (EPI) cells that have intercalated into the visceral endoderm (VE). ExE, extra-embryonic ectoderm-Scale bars: 50μm in low magnification images, 20μm in high magnification images. A, anterior; D, distal; P, posterior; Pr, proximal.

To infer directionality for the cross-over between the EPI and VE, we applied Palantir to Harmony augmented E3.5-E5.5 data after excluding TE/ExE cells (**Fig. 3b**). The same Nanog^hi^ cell as previously (**Fig. 2d**) was used as the starting cell. As expected, Palantir ordered cells along their developmental trajectories with high differentiation potential at E3.5, corresponding to the EPI-vs.-VE lineage divergence (**Fig. 3b**). In addition, an increase in differentiation potential was observed in a subset of EPI cells at the interface between EPI and VE at E5.5 (**Fig. 3b, black arrowhead**). This was followed by increasing VE branch probability (and concordant decrease in EPI branch probability) once the boundary of EPI cell density was crossed, continuing along the cells connecting the EPI and VE (**Fig. 3b, dotted lines**), suggesting that the observed continuity between these two lineages might result from a subset of EPI cells transdifferentiating into VE. Gene expression patterns of key anterior visceral endoderm (AVE), markers such as Lefty1, Cer1, Hhex ^49–51^, suggested that these EPI cells may contribute to AVE, a specialized cellular cohort exhibiting an intra-epithelial distal-to-anterior migratory behavior between E5.5 and E6.0 (**Fig. 3c**)^52^.

We used two *in vivo* lineage-tracing approaches to investigate cross-over and directionality between VE and EPI, two lineages believed to be mutually-exclusive. First, we crossed the EPI-specific *Sox2-Cre* and VE-specific *Ttr-Cre* mouse lines to the *ROSA26^mTmG^* reporter (**Fig. 3d**). Confocal imaging of E5.5-E6.0 *Sox2-Cre^TG/^+; ROSA26^mTmG/^+* embryos immunostained with a VE-specific anti-Gata6 antibody revealed the majority of Cre-recombined GFP-positive cells are located within the EPI. However, single GFP-positive cells were also observed within the emVE (yellow arrowheads, n=10/20 embryos analyzed), but not exVE. These trans-migrating cells were Gata6-positive, indicating they have trans-differentiated and acquired a VE identity. In E5.5 embryos, GFP-positive Gata6-positive cells were observed in distal locations, in the vicinity of the presumptive AVE, whereas by E6.0, they predominantly resided more anteriorly (**Fig. 3d, bottom**). By contrast, in *Ttr-Cre^TG/^+; ROSA26^mTmG^* embryos (n=0/27), all GFP-positive cells were restricted to the VE. These data support an EPI-to-VE trans-migration and trans-differentiation, but not vice-versa (**Fig. 3e**). To further validate these observations, we generated tetraploid embryo <-> *CAG-H2B-tdTomato* embryonic stem cell (ESC) chimeras, recovered embryos at E5.5-E6.0, and observed tdTomato-positive cells distributed throughout the EPI, and sparsely within the emVE, supporting an EPI-to-VE differentiation (n=19 analyzed, **Extended Data Fig. 7d**). By investigating gene cluster trends within the VE (**Extended Data Fig. 8a**) and EPI (**Extended Data Fig. 8b**) after excluding trans-differentiating cells (**Extended Data Fig. 8c**), we identified genes correlating with known endoderm factors such as Foxa2, Gata4, Gata6, Sox7 and Sox17, and pluripotency-associated factors such Nanog, Pou3f1 and Klf4 (**Supp. Table. 3**). Thus, trans-differentiation from EPI to VE, as predicted by Palantir, is verified by two independent lineage tracing approaches.

### Emergence of spatial pattern within the visceral endoderm epithelium at E5.5

By early post-implantation (E5.5, **Fig. 1**) the mouse embryo is radially symmetrical around a proximal-distal axis. The symmetry is broken, and the anterior-posterior axis is established through the anterior-ward migration of distal/anterior visceral endoderm (D/AVE) ^53^. Proximal-distal spatial patterning across the VE has been described preceding, and coincident to, the onset of gastrulation at E6.5, with a clear distinction between the morphology and function of the proximally located exVE (a cuboidal epithelium, overlying the ExE, giving rise to the yolk sac endoderm, YsE) and distal emVE (a squamous epithelium, overlying the epiblast, predominantly contributing to the gut endoderm). However, a day earlier, at E5.5, the VE appears morphologically uniform along the proximal-distal axis of the embryo^4,54,55,56^. To determine the onset of spatial patterning within the VE epithelium, we sought to establish when cells specified towards YsE-vs-gut tube could first be identified. We used Harmony to generate a unified view of cells of the VE lineage from E3.5 to E8.75 (**Fig. 4a**), and applied Palantir using a Nanog^hi^ ICM cell to define the start, with YsE and anterior/posterior ends of the gut tube as terminal states (**Fig. 4b, Extended Data Fig. 9a-c**). Since we ascertained that the probabilities towards anterior and posterior ends of the gut tube did not change until E7.5, we combined them to represent the probability of a cell committing to a gut tube fate (**Extended Data Fig. 9b-c**).

**Figure 4:**
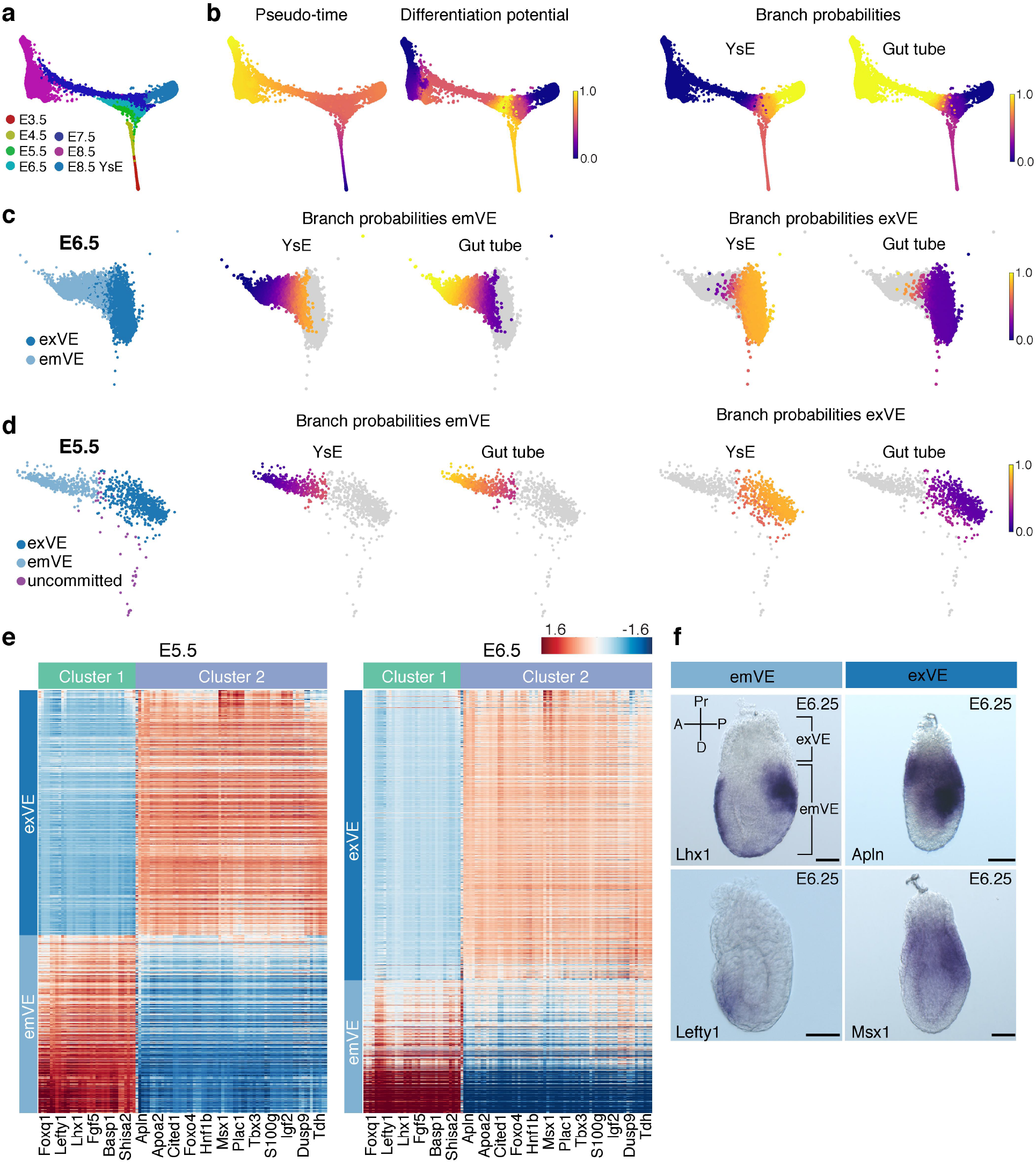
Emergence of spatial patterning within the visceral endoderm at the onset of post-implantation development (E5.5). The results shown were generated by pooling cells from all replicates from the E3.5-E8.75 time points and following Harmony augmentation. **a,** Force directed layout of trajectory of endoderm cells from blastocyst to midgestation, corresponding to E3.5-E8.75. **b,** Plots showing Palantir pseudo-time, differentiation potential and branch probabilities of endoderm cells from stages E3.5-E8.75 derived using a Nanog^hi^ cell as the start. **c,** Plots showing the branch probabilities of exVE and emVE cells at E6.5, as they commit towards YsE (extra-embryonic membrane) or (embryonic) gut tube. Cells labeled as exVE and emVE based on expression of known markers (left), match expected Palantir branch probabilities (right). **d,** Branch probabilities of E5.5 cells in committing towards YsE and gut tube were used to infer the putative exVE and emVE cell types at E5.5. Cells labeled as exVE and emVE based on Palantir’s branch probabilities. **e,** Heatmaps of genes expressed specifically in exVE or emVE at E5.5 (left). Cells are sorted within each compartment by pseudo-time ordering (left). These genes also distinguish exVE and emVE cells at E6.5 (right). (F) WISH of E6.25 embryos showing expression of Lhx1 and Lefty1, genes specific for emVE, and Apln and Msx1 specific for exVE. Scale bars: 50μm. A, anterior; D, distal; P, posterior; Pr, proximal.

The differentiation potential of cells at E3.5 and E4.5 remained unaltered suggesting the absence of specification towards YsE or gut tube, and hence an absence of spatial patterning at these stages (**Fig. 4b, Extended Data Fig. 9a and c**). By contrast, a clear distinction is evident between cells specifying towards YsE-vs-gut tube at E6.5 and E7.5 (**Fig. 4c, Extended Data Fig. 9e**). Based on the expression of marker genes in scRNA-seq data, and correlation to bulk RNA-seq gene expression of sorted E7.5 exVE and emVE tissues, we determined that cells specifying towards YsE and gut tube are exVE and emVE, respectively, consistent with reported spatial patterning at these stages (**Extended Data Fig. 9d**). Interestingly, VE cells at E5.5 resemble both earlier and later stages: a subset did not exhibit any change in differentiation potential indicating a more uncommitted state (**Extended Data Fig. 9f**), while the majority exhibited an altered differentiation potential indicating propensity towards YsE-vs-gut tube (**Fig. 4d, Extended Data Fig. 9f**). Taken together, these data reveal that, at the transcriptional level, spatial patterning can be detected at E5.5, preceding the onset of morphological changes within the VE epithelium.

### exVE is a uniform cell type, while emVE comprises two types of cells, AVE and non-AVE

We reasoned that the spatial patterning would be accompanied by coordinated changes in gene expression programs conferring exVE/emVE identity. Differential expression between bulk RNA sequenced sorted exVE and emVE populations at E7.5 revealed substantially more upregulated genes in emVE compared to exVE (2239 genes in emVE *vs*. 291 genes in exVE, (**Extended Data Fig. 9d**). This suggests that emVE represents a specialized variant of exVE, perhaps activating a gene expression program in response to stimuli, such as BMP or Nodal^57–60^. To explore this further, we computed covariance matrices of highly variable genes for putative E5.5 exVE and emVE populations (**Extended Data Fig. 9g**). Clustering of the exVE covariance matrix did not reveal any coherent clusters, but clustering of the emVE covariance matrix uncovered two clusters, with genes within each cluster being highly correlated, but strongly anticorrelated with genes of the other cluster (**Extended Data Fig. 9g**), consistent with the observations from bulk data that suggested emVE cells representing a specialized exVE subpopulation. The co-varying genes not only exhibited contrasting expression patterns in putative E5.5 exVE and emVE cells (**Fig. 4e**) but can also distinguished *bona fide* exVE and emVE cells at E6.5 and E7.5 (**Fig. 4e, Extended Data Fig. 9h**). This indicates that these genes comprise a core transcriptional program conferring exVE and emVE identity at E5.5, preceding the appearance of morphological changes. emVE specific genes (**Fig. 4e, Cluster 1**) included Lhx1 and Lefty1, and other AVE-specific genes such as Cer1 and Hhex^49,51,53^. exVE specific genes (**Fig. 4e, Cluster 2**) included Apln and Msx1 (**Fig. 4e, Extended Data Fig. 9i, Supp. Table. 4**). ISH for Lhx1, Lefty1 and Apln, Msx1 on E6.25 embryos validated em/exVE regionalized expression (**Fig. 4f**). In sum, these data demonstrate that the visceral endoderm epithelium is patterned at the very earliest stages of post-implantation development, and that emVE represents a derivative identity of exVE, also encompassing the AVE subpopulation.

### AP patterning occurs concomitant with the formation of the gut tube

We next investigated the convergence of visceral (VE) and epiblast-derived definitive endoderm (DE) cells within the emergent gut endoderm and explored the spatial distribution of their descendants along the anterior-posterior (AP) axis of the gut tube. We pooled data from the anterior/posterior E8.75 gut tube compartments with the (Afp-)GFP-positive/negative sorted populations after correcting for batch effects (**Fig. 5a**). To generate a unified view of all gut tube cells, we used a manifold classifier based on Palantir to infer the GFP status of the anterior/posterior cells, and the AP position of the GFP-positive/negative cells (**Extended Data Fig. 10a and b**). The strongest signal in the data, as determined by the first diffusion component, was the ordering of cells along the AP axis (**Extended Data Fig. 10c**). To confirm that the AP ordering of cells reflected their spatial distribution along the gut tube, we compared gene expression from scRNA-seq data, ordered based on the first diffusion component, with bulk RNA -seq data of micro-dissected quadrants of gut tube and determined that Nkx2-1, an anterior gene and Hoxb9, a posterior gene show consistent expression patterns (**Extended Data Fig. 10c**). To determine a more robust ordering rather than relying on a single diffusion component, we determined an estimate of the pseudo-spatial ordering of cells along the gut tube by computing multi-scale distances from the anterior-most cell after projecting the cells onto diffusion components (**Fig. 5b, Extended. Data. Fig. 10d**). We then binned cells along the AP pseudo-space ordering and computed the proportion of VE and DE descendants in each bin to reveal that VE and DE descendants exhibited distinct distributions along the AP axis. As expected from our previous studies ^9^, we observe extensive intermixing of VE and DE descendants along the AP pseudo-space axis with enrichment of DE descendants in the anterior, and VE descendants in the posterior gut tube (**Fig. 5b**). To determine if VE cells, being of extra-embryonic origin, attained a level of equivalence to DE descendants, we compared the expression of markers of the primordial endodermal organs within the two populations: Nkx2-1 (thyroid/thymus)^61^, Irx1 (lung)^62^, Ppy (liver)^63^, Pdx1 (pancreas)^64^, Fabp1 (small intestine)^65^ and Hoxb9 (posterior). With the exception of Nkx2-1, which is expressed in the anterior-most cells of the gut tube and is therefore exclusive to DE, all other genes were expressed at significant levels in both VE and DE cells, and at similar AP positions (**Fig. 5c**). Furthermore, we noted a strong correlation in global gene expression patterns between VE and DE cells in bins along the AP pseudo-space (**Fig. 5b, purple**) demonstrating that VE and DE descendants become similarly patterned and acquire regionalized organ-specific identities, each likely contributing descendants to endodermal organs (**Fig. 5c**).

**Figure 5:**
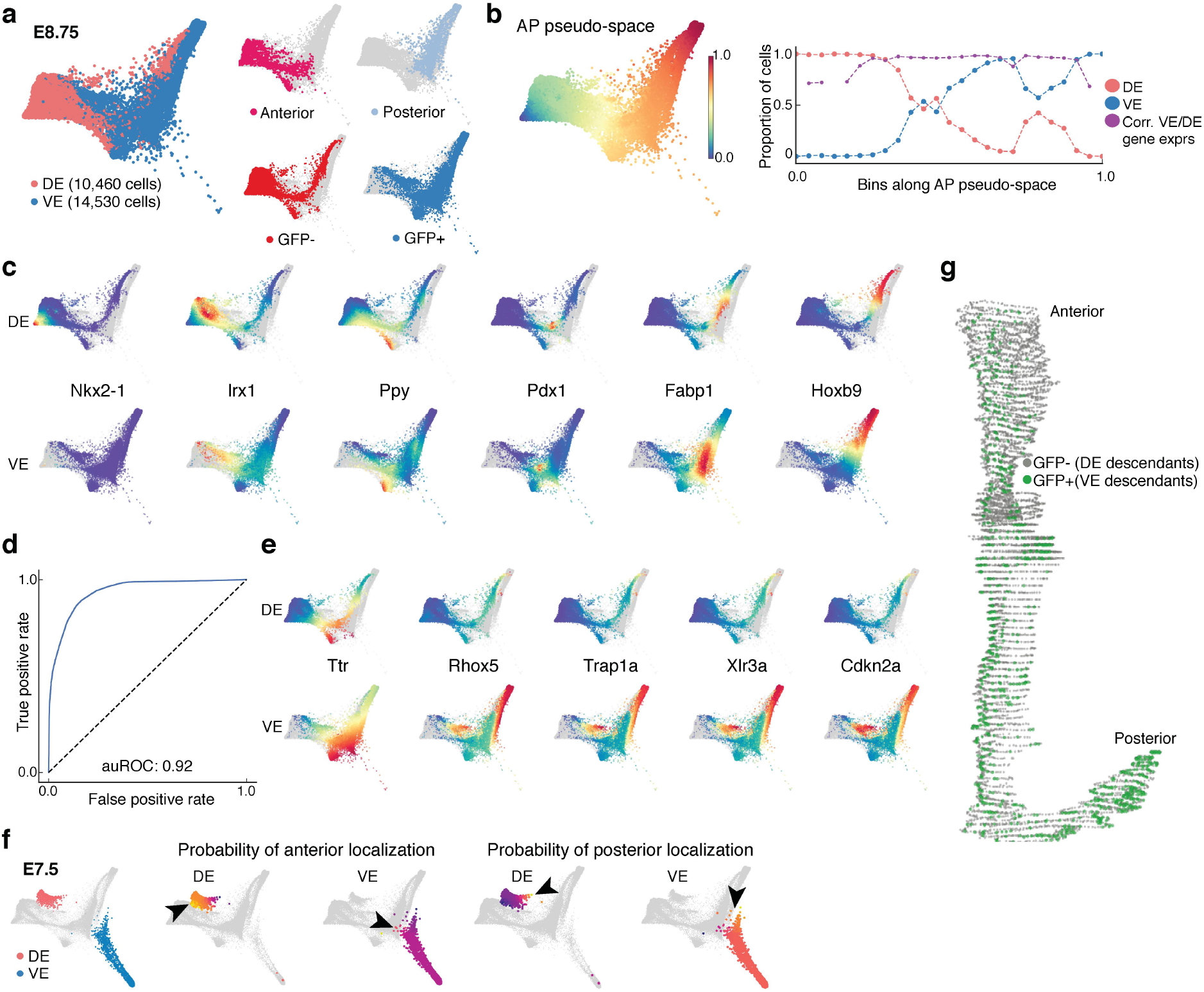
Anterior-Posterior pseudo-spatial axis of cells residing within the E8.75 gut tube. Force directed layout of the VE and DE cells at E8.75 derived by combining the anterior/posterior cells with the Afp-GFP-positive/AFP-GFP-negative sorted cells using mNNCorrect batch correction ^27^(panels A-C,E). **a,** Cells colored based on measured or inferred AP position (left), cells labeled by measured data (right). **b,** Plot showing the inferred anterior-posterior (AP) pseudo-space ordering of the E8.75 cells (left) and the proportion of VE and DE cells in bins along the AP pseudo-space ordering (right). Purple dots represent the correlation of aggregate expression VE and DE cells in each corresponding bin. **c,** Plots showing the expression patterns of key organogenesis markers in the DE cells (top) and VE cells (bottom). **d,** Receiver operating curve for classification of E8.75 VE and DE cells using the model trained with the E7.5 VE and DE cells. **e,** Plots showing the expression patterns of genes (top - DE; bottom -VE) that are best predictive of the VE class in the VE-DE classifier. **f,** Force directed layouts following Harmony of E7.5 and E8.75 VE and DE cells with the E7.5 cells highlighted (left). E7.5 VE and DE cells colored by the branch probability of anterior localization (middle) and posterior localization (right). Black arrow heads indicate the early emergence of AP spatial patterning at E7.5 with E7.5 DE cells predominantly destined for anterior location whereas E7.5 VE cells predominantly destined towards the posterior. **g,** A 3D rendering of the gut tube showing all cells along the A-P axis. Nuclei of VE and DE cells are labeled in green and grey, respectively.

### VE cells possess a memory of their lineage history

Despite this global similarity in transcriptomes, we explored whether VE and DE descendants retain a transcriptional memory of their lineage history. To overcome the confounding effects introduced by the spatial distribution of VE and DE descendants along the gut tube, we trained a sparse logistic regression model to classify E7.5 VE and DE cells using all genes as features. This classifier achieved near-perfect accuracy (auROC:0.96), demonstrating that at E7.5, VE and DE cells are transcriptionally distinct (**Extended Data Fig. 10e**), and indeed occupy distinct regions in the force-directed map (**Fig. 5f, right**). We next applied this classifier (trained on E7.5) to predict the origin of cells within the E8.75 gut tube, achieving a similarly high accuracy (**Fig. 5d,** auROC: 0.92). Thus, demonstrating that despite extensive morphological (e.g. gut endoderm becoming internalized) and transcriptional changes taking place between E7.5 and E8.75, lineage history of VE and DE descendants is maintained by means of a core subset of genes. Genes such as Rhox5, Trap1a, Xlr3a, Cdkn2a and Ttr are predictive of the VE identity (**Fig. 5e**), whereas ribosomal genes Rplp1, Rps23, Rpl11 and Rplp0, as well as Nnat are predictive of DE identity (**Extended Data Fig. 10f, Supp. Table. 5**). Notably, of the VE-specific genes, Rhox5^66^ and Xlr3a^67^, are X-linked.

### Emergence of spatial distribution of VE and DE cells along the gut tube at E7.5

Since the classifier trained on E7.5 cells could accurately identify DE and VE cells within the E8.75 gut tube, we sought to determine whether spatial patterning of cells along the AP axis could be observed earlier. We applied Palantir separately for DE and VE cells after MNN augmentation using E7.5 GFP-negative and E8.75 GFP-negative cells for DE, and E7.5 GFP-positive emVE and E8.75 GFP-positive cells, respectively, using anterior and posterior cells as the terminal states. Our results revealed the presence of a small fraction of cells already acquiring AP identities in both VE and DE compartments at E7.5 (**Fig. 5f**). Notably, the DE cells were predominantly primed towards an anterior localization, whereas VE cells were predominantly posteriorly (black arrowheads in **Fig. 5f, Extended Data Fig. 10g**). We next compared the distribution of VE cells along the AP axis of embryos, with VE proportions inferred from the scRNA-seq data along the pseudo-space axis. To quantify the distribution of VE/DE cells within gut tubes of E8.75 (13ss) embryos at cellular resolution, complete serial transverse sections of three Afp-GFP^TG/+^ embryos were analyzed; all endoderm cells counted and assigned a VE (GFP-positive) or DE (GFP-negative) identity (**Supp. Table 12**). A representative 3D reconstruction of the entire gut tube from one of the embryos analyzed was generated using Neurolucida software (**Fig. 5g,** green/VE nuclei, grey/DE nuclei). Comparison of the VE descendant proportions in bins along the AP axis of the gut tube, and the AP pseudo-space axis highly correlated (**Extended Data Fig. 10h**), further demonstrating the accuracy of the inferred AP pseudo-space. The pseudo-space ordering is also robust to different parameters and reproducible across replicates (**Extended Data Fig. 10i-k**).

### Emergence of organ identities across the AP axis of the gut tube

The gut tube comprises progenitors of the endodermal organs, which will later on emerge in a stereotypical AP sequence: thymus, thyroid, lung, stomach, liver, pancreas, small intestine and colon. To investigate whether the E8.75 gut tube already contains information relating to the later establishment of endodermal organs, we clustered all gut tube cell populations (anterior /posterior, GFP-positive/GFP-negative) and annotated clusters based on differential expression of primordial organ markers. We then determined an ordering of these clusters along AP pseudo-space by computing the average distance from the anterior most cell. The resulting ordering of clusters in AP pseudo-space perfectly matched the order in which organ identities appear along the AP axis of the gut tube (**Fig. 6a**). We observed a high degree of variability in the density of cells along the pseudo-spatial axis (**Fig. 6b and Extended Data Fig. 11a-b**), with low density regions between clusters, potentially demarcating distinct organ precursors. As expected, the proportion of VE cells contributing to each cluster increased with posterior localization of clusters, with most VE cells contributing to the prospective large intestine/colon (**Fig. 6c**). Cluster-specific expression patterns were validated using WISH on gut tubes, confirming the accuracy of inferred AP pseudo-space, and the emergence of endodermal organ precursors at E8.75 (**Extended Data Fig. 11a**).

**Figure 6:**
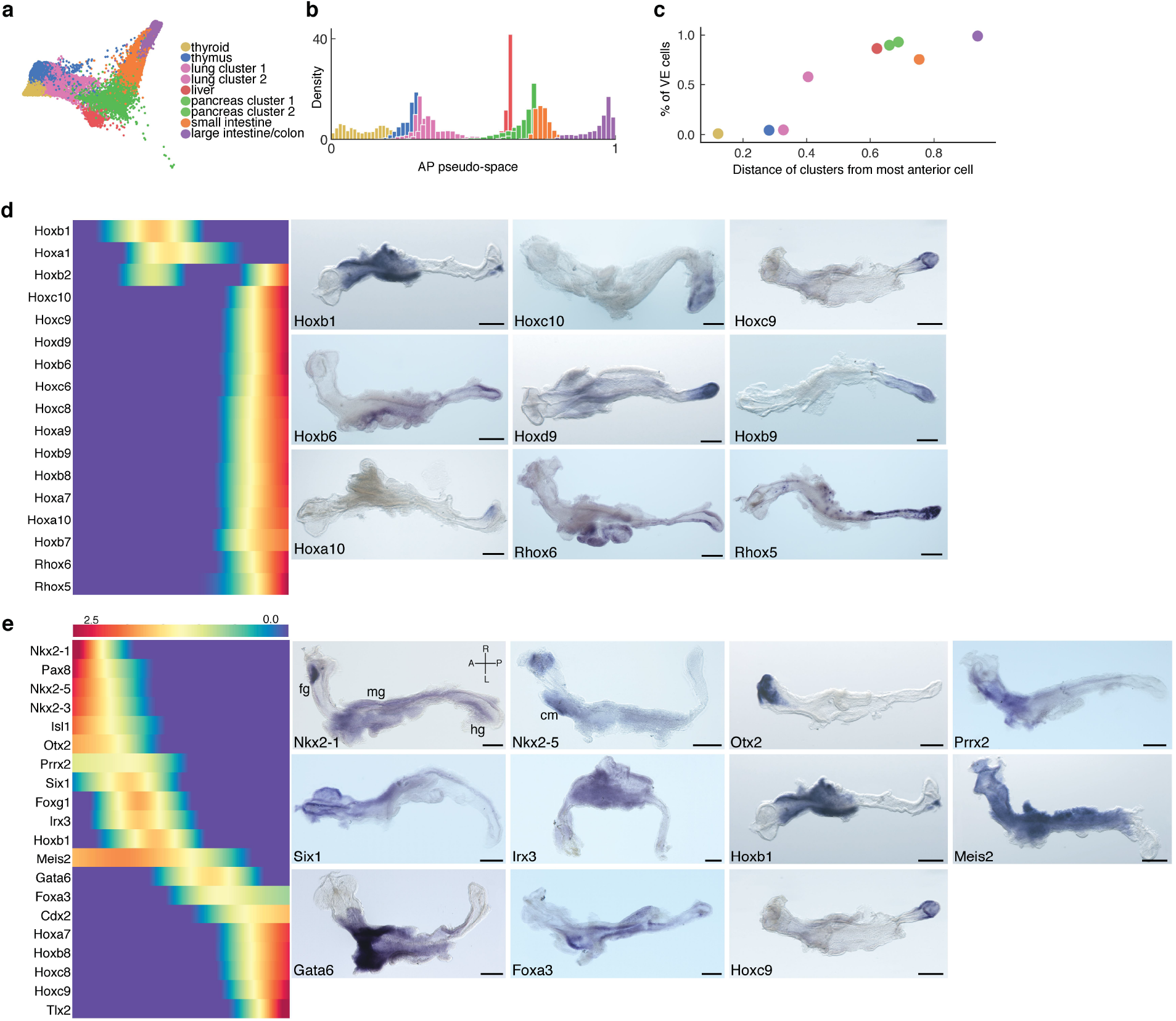
Spatial patterning of the E8.75 gut tube. **a,** Force directed graph of E8.75 cells colored by Phenograph clusters annotated with the putative primordial endodermal organ associated with each cluster. **b,** Plot showing the density of cells of the different clusters along the AP pseudo-space axis, each distribution represents cells from one Phenograph cluster. **c,** Plot showing the percentage VE cells in each cluster, where the clusters are ordered by average distance from the anterior trip of the AP pseudospace axis. **d,** Heatmap showing the expression patterns of the Hox family genes along the AP pseudo-space axis. Right: Validation of Hox gene expression patterns by WISH on gut tubes micro-dissected from E8.75 embryos. **e,** Expression patterns of TFs most predictive of the AP pseudo-space in a regression model. Columns represent cells ordered by pseudo-space and each row represents the expression of a particular TF. TFs were ordered based on their expression along the AP axis. Validation of predictive A-P expression patterns by WISH of TFs on gut tubes. All scale bars: 200μm, except for Nkx2-1, Irx3: 100μm. A, anterior; cm, cardiac mesoderm; fg, foregut; hg, hindgut; L, left; mg, midgut; R, right; P, posterior. All scale bars: 200μm, except for Hoxc10: 100μm.

We next sought to identify the determinants of AP patterning in the gut tube. Hox genes which are colinearly expressed, at least in the developing central nervous system, are considered canonical descriptors of position along the AP axis^68,69^. Surprisingly, we found that the majority of Hox genes expressed within the E8.75 gut tube to be predominantly posteriorly localized (**Fig. 6d and Extended Data Fig. 12**), suggesting that AP identity within the gut tube might be independent of a Hox code. Furthermore, WISH of gut tubes demonstrated that the domains of Hox gene expression were not always correlated between the endoderm and surrounding tissues (**Extended Data Fig. 12:** Hoxb1, Hoxd4).

Given that Hox genes did not appear to be the determinants of AP patterning in the gut tube, we next explored whether cell-autonomous signals through transcriptional regulation might contribute to AP patterning. We trained a sparse regression model to predict AP pseudo-space (**Fig. 4**) using the expression of all transcription factors (TF) as features. This revealed TF expression was exceptionally accurate in predicting AP pseudo-space order (**Extended Data Fig. 11c,** Correlation: 0.97), strongly indicating that transcriptional regulation, possibly in response to signals from the mesoderm, plays a key role in establishing the AP pattern of the gut tube. The sparse regression model also allowed identification of a core group of TFs sufficient for predicting the AP pseudo-space axis (**Fig. 6e and Extended Data Fig. 11d**). The TFs and their coefficients are extremely robust to sub-sampling of data further demonstrating their predictive power (**Extended Data Fig. 11e-f**). As expected, the core set of factors showed strong spatial patterns of expression, validated using WISH; from Nkx2-1 at the anterior, to Hoxb9 and Hoxc9 at the posterior (**Fig.6**).

## DISCUSSION

Single-cell transcriptomic analyses have revealed unappreciated complexities and dynamics encompassing cell fate decisions during developmental progression^19,22,24,25,70^. To understand the ontogeny of the endodermal organs in mammals, we analyzed 112,217 single-cell transcriptomes (**Fig. 1**) representing the pre-implantation blastocyst (emergence of the first endoderm population) to midgestation (formation of the gut tube). Combining Palantir^6^ with our new Harmony algorithm empowered us to computationally organize 112,217 endodermal sc-transcriptomes based on pseudo-time and pseudo-space, identifying unappreciated lineage relationships and the emergence of a stereotypical patterned organization within the gut tube.

Our high-resolution transcriptomic data reveal an early heterogeneity within the ICM, which combined with Palantir’s analysis elucidated the detailed structure of the ensuing cell fate decision towards PrE or epiblast. Our analysis, could pinpoint the order and timing of events during development, including the expression trends of key developmental regulators, such as FGF signaling pathway components. For example, we could observe that Fgrf1 and Fgrf2 are coordinately expressed during a very narrow time-window at E3.5 while PrE is being specified. By E4.5, both Fgfr3 and Fgfr4 are expressed in PrE, while Fgfr1 and Fgfr2 are downregulated, suggesting that an FGF signal may be reiteratively used, but differentially transduced within a single lineage.

Our analysis reveals that throughout embryogenesis cells acquire a transcriptional identity specifying their future cell fate and spatial positioning well before cells morphologically change and migrate to their predetermined (based on transcriptional identity) position. At 3.5, the morphologically similar ICM already transcriptionally distinguishes two sequential uncommitted states, as well as cells having acquired an EPI or PrE identity. At E5.5, before proximal-distal symmetry is broken, cells already begin to acquire transcriptional identities of the proximal exVE and distal emVE. Similarly, we already see transcriptional priming of spatial patterning of cells along the Anterior-Posterior (AP) axis at E7.5. By E8.75, the top diffusion component, which represents the strongest axis of variation in the data, recapitulates pseudo-space directly from the transcriptomic information, without any spatial alignment or landmarks. Along the AP axis of the gut tube, we already observe organ precursors, expressing organ specificities in their stereotypical order along the gut tube.

While cells developed a marked propensity towards specific cell fates earlier than previously appreciated, they nevertheless retain a remarkable degree of plasticity. Most notably, after EPI and PrE/VE diverge into distinct lineages at E3.5, we observe a trans-differentiation event between these two very distinct lineages. While the PrE/VE and EPI are considered lineage-distinct, they lie in close apposition to one another as embryos develop from the late blastocyst to early post-implantation (E5.5). Application of Palantir to our data revealed an unprecedented lineage relationship between EPI and VE, whereby EPI cells trans-migrate into the VE epithelium, and concomitantly trans-differentiate, acquiring a VE identity. We validated Palantir’s computational prediction with lineage-tracing experiments in embryos, supporting the unidirectional EPI-to-VE translocation, and concomitant trans-differentiation, demonstrating EPI descendants actively contribute to the VE (**Fig. 3**). It is unclear whether this EPI-to-VE differentiation reflects a removal of ‘less-fit’ cells from the pluripotent epiblast compartment ^72–75^ or an active recruitment of cells to the VE. In the context of cell competition-based model for the EPI, cell engulfment has been proposed as the mechanism of cell removal. In support of the active recruitment of EPI cells to VE, it has been previously suggested that breaks in the basement membrane separating the EPI from VE, might allow cells to escape the EPI layer, and populate the nascent DVE^76^.

A second notable event occurs between E7.0 and E7.5, when EPI-derived (embryonic) definitive endoderm (DE) intercalates with the emVE to form the gut endoderm, an epithelium comprising cells of two distinct origins^7,9^. By the time emergent gut endoderm has become internalized to form a gut tube (E8.75), we could identify clusters of cells expressing markers of organ identity and correlating in AP pseudo-space with the stereotypical order of the emergent endodermal organs (**Fig. 6**). Importantly, while VE and DE retain some small signature of their lineage history, they largely acquire transcriptomic equivalence. Expression patterns are largely determined by spatial localization along the AP, rather than VE/DE lineage history.

Our data suggests that cell fate is determined through a combination of a cell-intrinsic propensity to specific fates and extrinsic signaling from the environment. At each stage we found evidence of cells molecularly poised for specific spatially patterned fates, well before overt spatial organization. On the other hand, we noted that expression patterns are determined based on their spatial position within the gut tube rather than their origin, likely due to cell-cell signaling. It will be interesting to dissect the interplay between intrinsic and extrinsic factors in determining cell fate decisions, using quantitative imaging, epigenetic data and genetic perturbations.

## Acknowledgements

We thank S.Y. Kim at the NYU Rodent Genetic Engineering Core Facility for generating tetraploid chimeras; L. Beccari, D. Duboule, M. Torres, L. Sussel, D. Wellik, M. Wilkinson for plasmids; J. Brickman for ESCs. This work was supported by grants from the NIH (R01-HD094868 and R01-DK084391 to A.K.H., and DP1-HD084071 and R01-CA164729 to D.P.), MSKCC Society for Special Projects (to A.K.H. and D.P.), and NSERC (RGPIN-2018-05018 to P.H.).

## Author information

### Author contributions

S.N., M.S., D.M.C., P.A.H., A.K.H. and D.P. conceived the project. S.N., Y.Y.K and V.G. and collected single cells from embryos with assistance from C.S.S., N.S. and A.K.H. S.N., Y.Y.K. and R.G. performed FACS of Afp-GFP cells. S.N., V.G. and S.C.B. generated libraries for scRNA-seq. S.C.B. and D.C. performed RNA-seq. M.S. and D.P. conceived and developed the Harmony and Palantir algorithms. M.S., V.L. and R.S. performed computational analyses of scRNA-seq and bulk RNA-seq data. V.G. performed IF staining of preimplantation embryos. S.N. and Y.Y.K. performed lineage-tracing, IF and WISH experiments on post-implantation embryos and gut tubes. S.N., M.S., A.K.H. and D.P. analyzed and interpreted the data, and wrote the manuscript with input from all authors.

### Competing interests

S.C.B and D.M.C. are employees and shareholders at 10x Genomics.

### Corresponding authors

Anna-Katerina Hadjantonakis & Dana Pe’er

### Data and software availability

All the generated data including bulk and scRNA-seq data will be made available through GEO. Harmony is available as a python module here: https://github.com/dpeerlab/Harmony. A Jupyter notebook detailing the usage of Harmony along with sample data is available here:http://nbviewer.jupyter.org/github/dpeerlab/Harmony/blob/master/notebooks/Harmony_sam_ple_notebook.ipynb

## Extended Data Figures

**Extended Data Figure 1:**
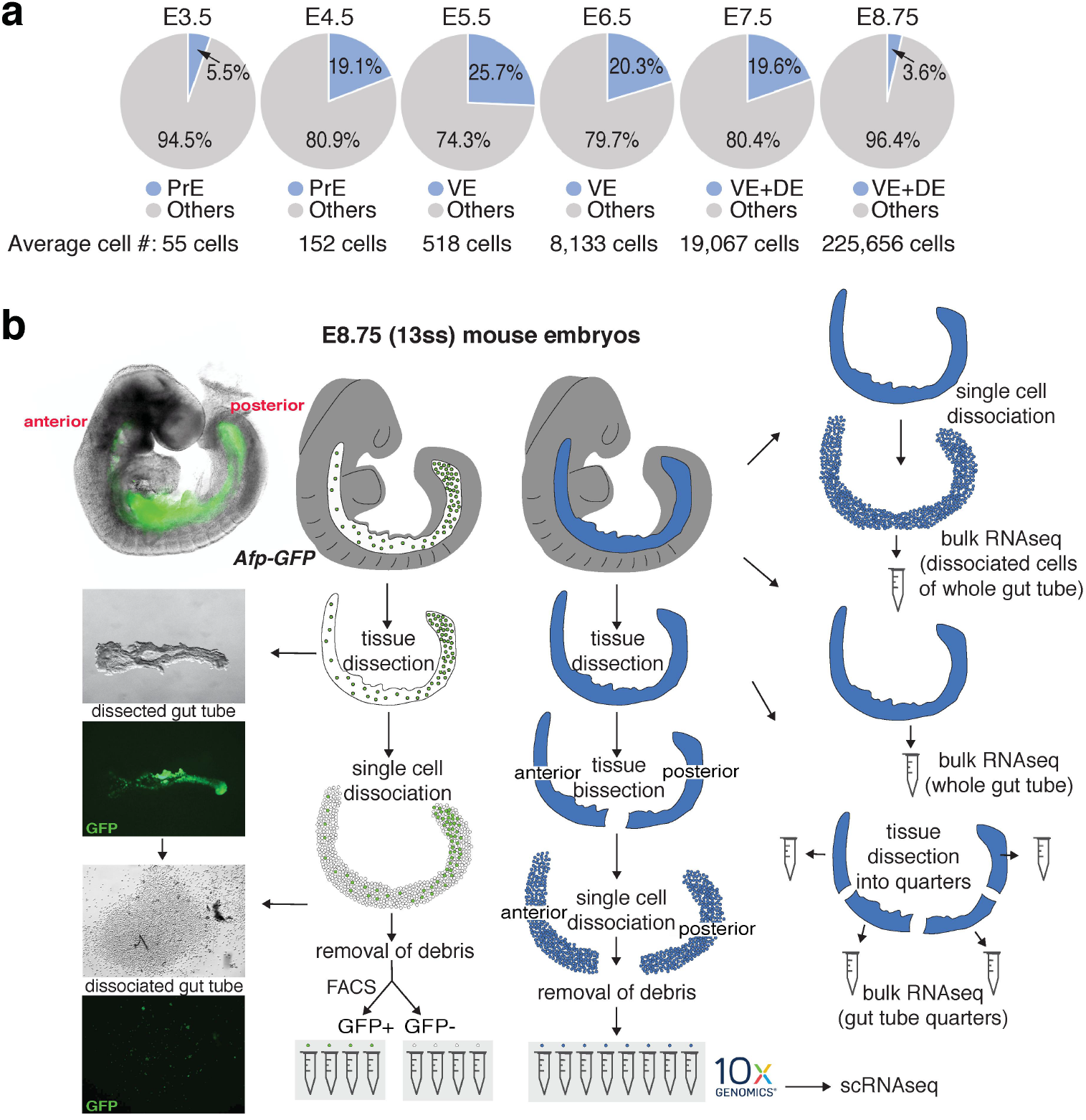
Endoderm cell representation in mouse embryos, from blastocyst through midgestation, and single-cell collection pipeline. **a,** Pie charts showing fraction of endoderm cells per embryo, for all stages used in study. **b,** Schematic of protocol used for single cell collection, with the E8.75 gut tube provided as an example. Gut tubes were micro-dissected from embryos and then dissociated into single cells. Single cells either of anterior and posterior halves of gut tubes, or AFP-GFP-positive (VE descendants) and AFP-GFP-negative (DE descendants) collected using FACS, were used for single cell library construction using the 10x Chromium platform. For bulk RNA-seq, whole gut tubes that had been dissociated into single cells and the pooled, whole gut tubes, and whole gut tubes dissected into quarters, were collected for sequencing.

**Extended Data Figure 2:**
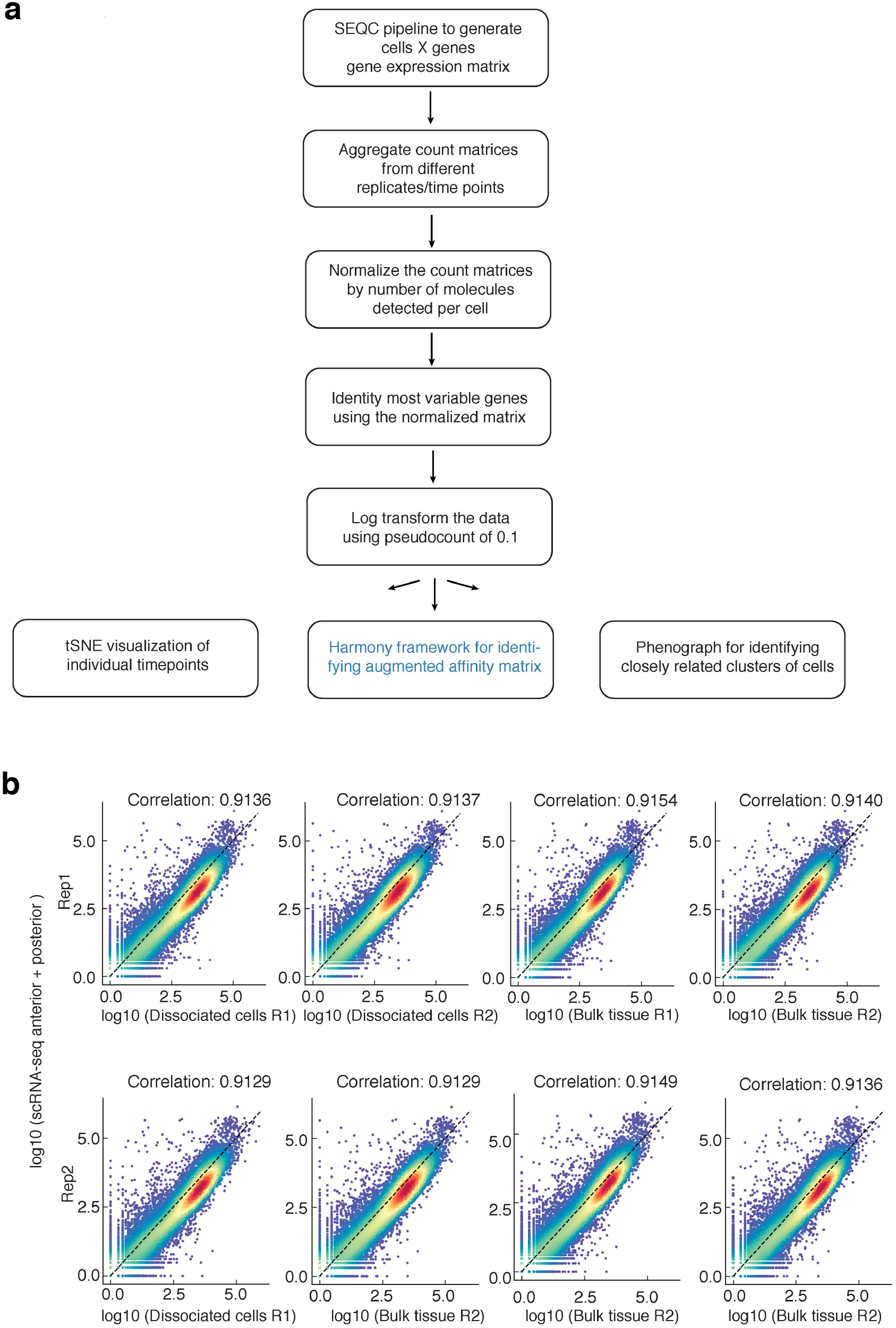
Computational pipeline and comparison of scRNA-seq with bulk RNA-seq data. **a,** Flow chart of computational pipeline for data processing. **b,** Plots showing the correlation between aggregated scRNA-seq data of anterior and posterior halves of the gut tube with bulk RNA-seq of dissociated cells and bulk tissue, respectively. The two rows represent the two replicates.

**Extended Data Figure 3:**
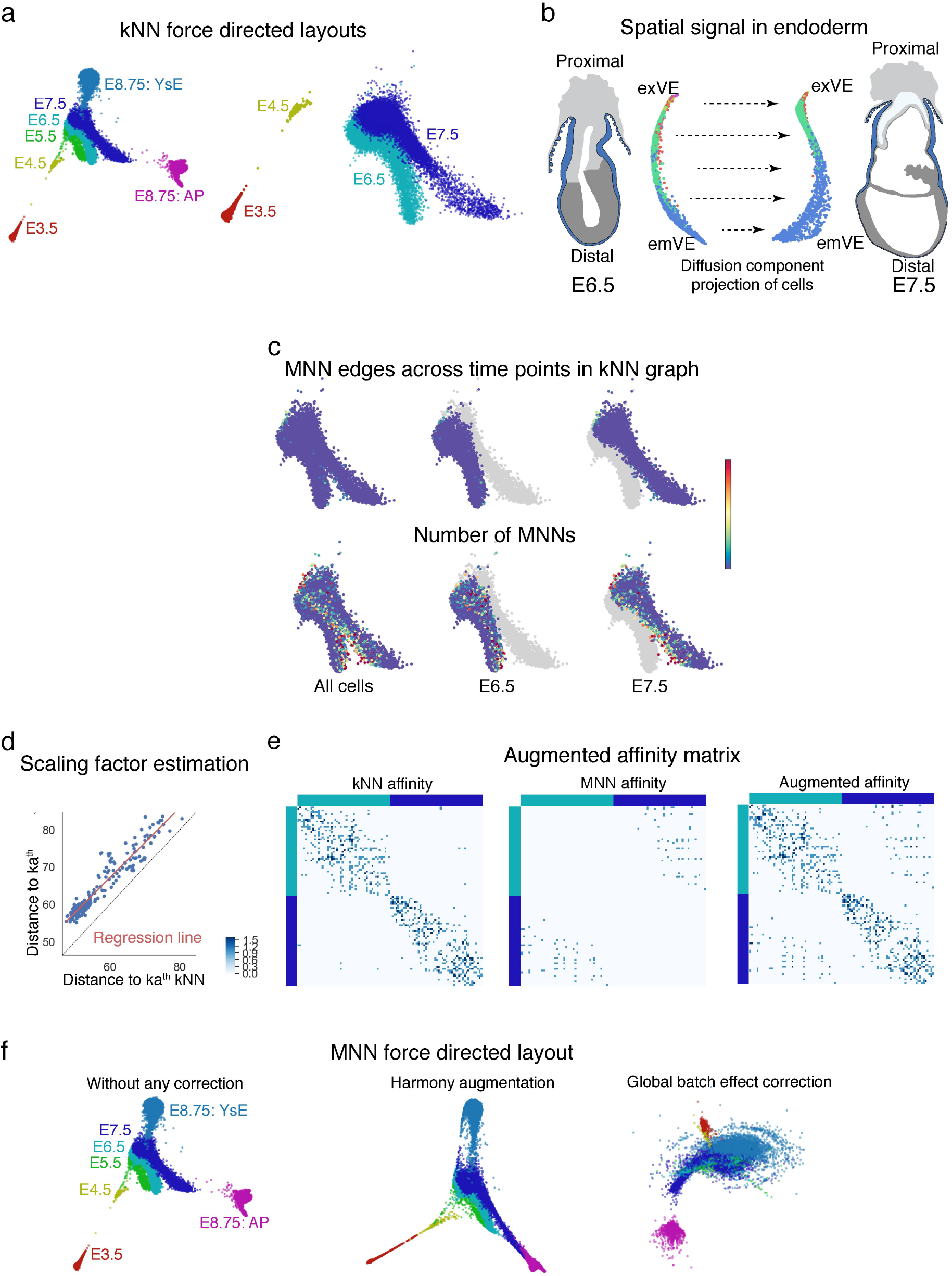
MNN augmentation to correct batch effects between time-points. **a,** Force directed layouts for cells from the following time-points: E3.5, E4.5, E5.5, E6.5, E7.5 and E8.75 (amalgamation of anterior and posterior halves). Cells are colored by time point. The graph was generated using an adjacency matrix derived from the kNN graph. Differences between consecutive time-points represent underlying developmental changes but are confounded by technical batch effects, including discontinuity between E3.5 and E.4.5 and lack of spatial alignment between E6.5 and E7.5. **b,** E6.5 and E7.5 cells projected along their respective first two diffusion components. These projections reveal the presence of strong spatial signal in the individual time-points. Cells are colored by Phenograph clusters. **c,** The number of edges connecting cells between time-points are limited in the kNN graph (Top panel). Bottom panel: Plots showing the number of mutually nearest neighbors (MNNs) between E6.5 and E7.5 time-points. The MNNs are enriched along the boundary between time points, supporting augmentation of the kNN graph with additional edges between mutually nearest neighbors (MNNs) between the consecutive time-points. d, The MNN distances can be converted to affinities on a similar scale as the kNN affinities, using linear regression to determine the relationship between the *ka^th^* kNN and *ka^th^* MNN distances. **e,** Example of the augmented MNN affinity matrix construction. Left panel: kNN affinities for a subset of E6.5 and E7.5 cells. Middle panel: MNN affinity matrix constructed using linear regression (d) to convert distances E6.5 and E7.5 cells to affinities. Right panel: Augmented affinity matrix: Sum of the kNN and MNN affinity matrices. **f,** Comparison of the force directed layouts. Left: Standard kNN affinity matrix, Middle: Harmony’s augmented affinity matrix. Right: Plot generated using mnnCorrect ^27^ for global batch effect correction which leads to “over-correction” and loss in signal between time-points.

**Extended Data Figure 4:**
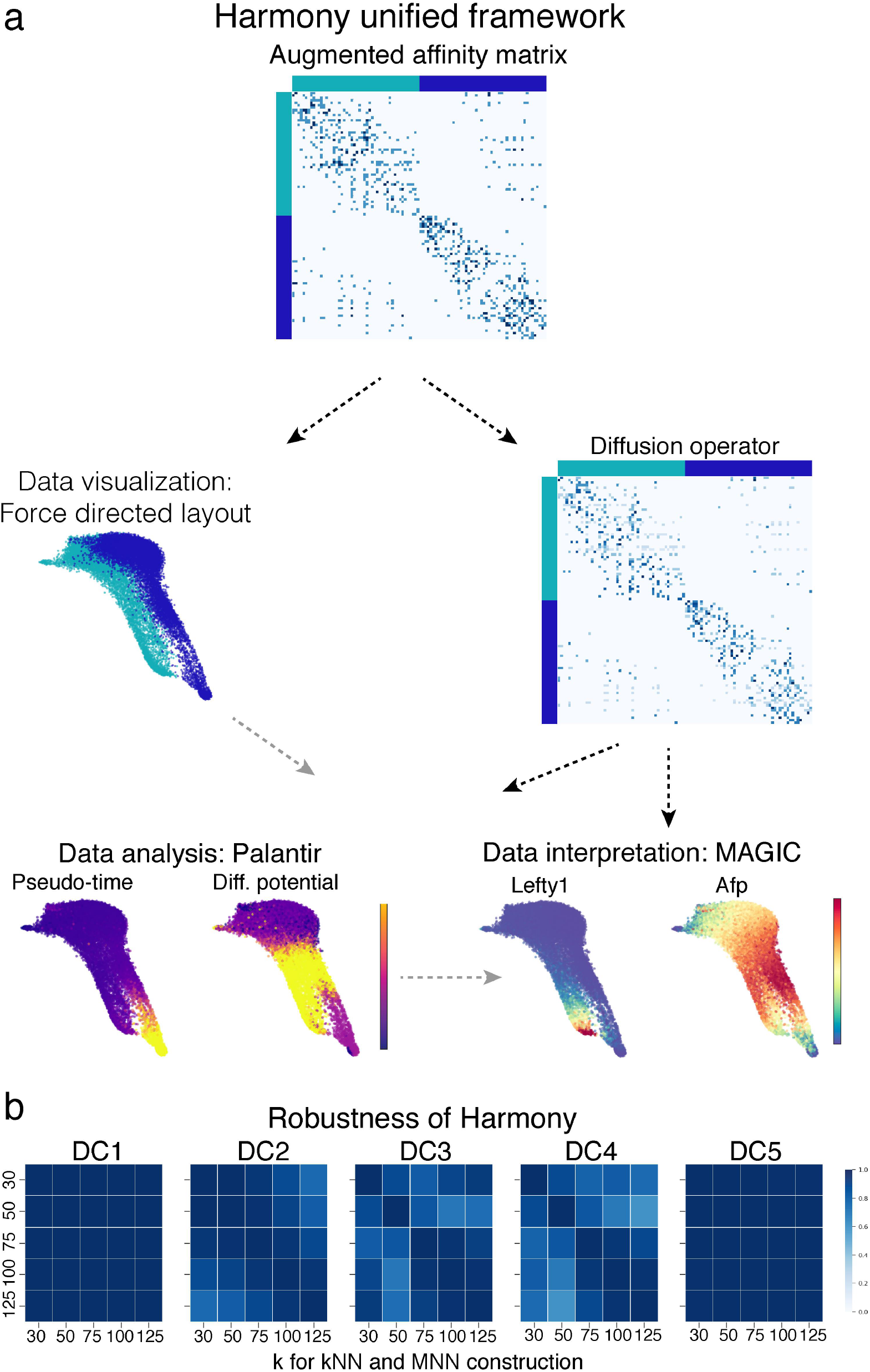
Harmony unified framework for scRNA-seq data analysis. **a,** Harmony framework starts with the augmented affinity matrix generated as shown in **Extended Data Figure 3** and described in supplemental methods. The augmented affinity matrix is used to generate the force directed graph for visualizing the data. The same augmented matrix is used to compute the diffusion operator for determining the diffusion components which (a) forms the basis for Palantir trajectory detection forming the data analysis arm, and (b) data interpretation by using MAGIC imputation. **b,** Robustness of Harmony: Plots showing the correlation between diffusion components for different values of k, the number of nearest neighbors for kNN graph construction. VE cells in **Fig.4** were used for testing robustness.

**Extended Data Figure 5:**
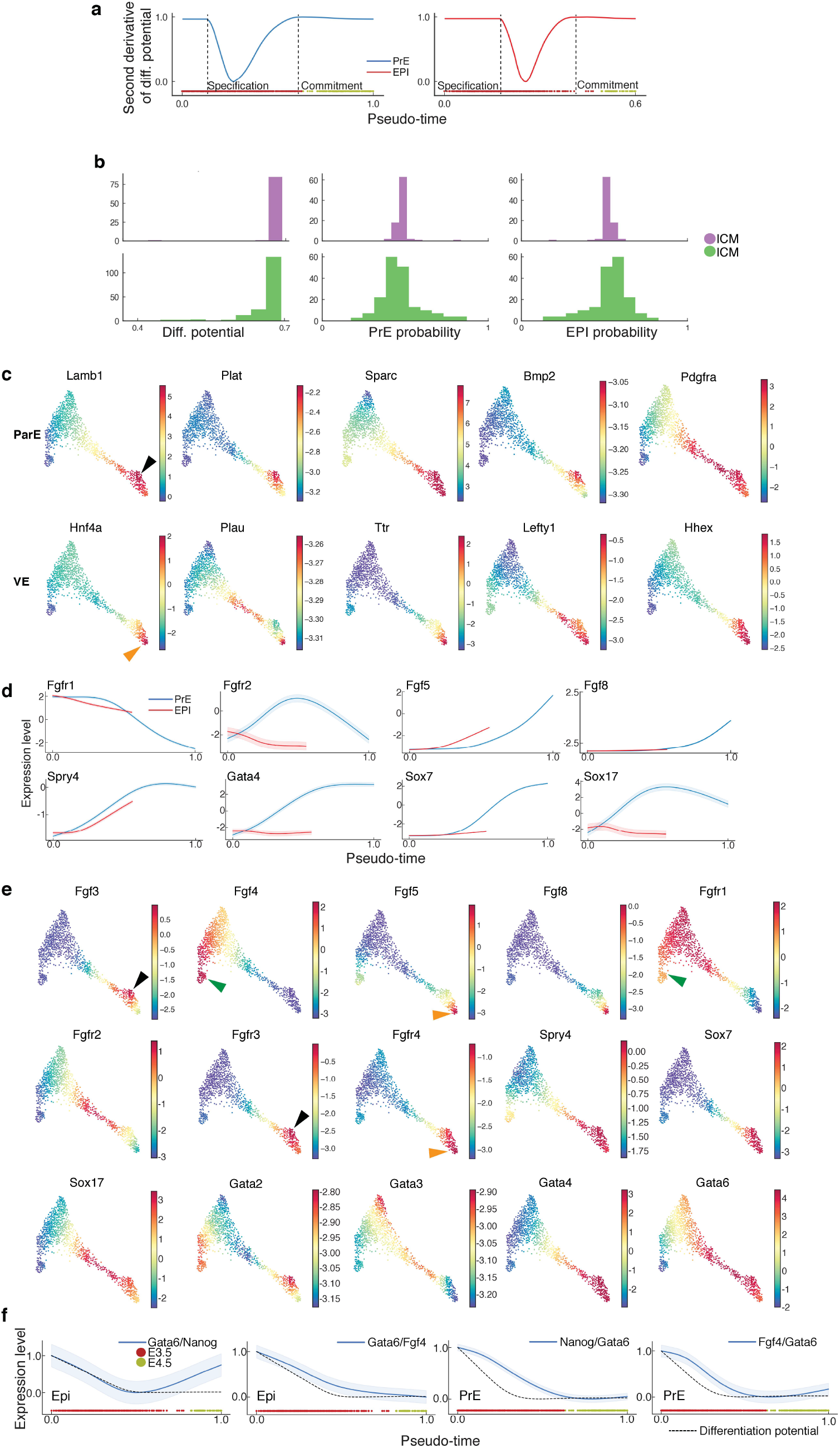
Gene expression trends in EPI, PrE, VE and ParE lineages in the blastocyst. **a,** Plots showing the second derivative of PrE and EPI differentiation potential along pseudotime demonstrating that changes in differentiation potential and hence lineage commitment in both lineages occur at E3.5. Points of highest changes along pseudo-time represent lineage specification and commitment. **b,** Histograms showing the distribution of differentiation potential (left), PrE fate probability (middle) and EPI fate probability (right) in the E3.5 ICM clusters. **c,** Gene expression patterns of parietal (ParE) and visceral endoderm (VE) markers. Each cell is colored based on its MAGIC ^30^ imputed expression level for the indicated gene. Black and orange arrowheads mark presumptive ParE and VE lineages, respectively. **d,** Plots comparing gene expression trends along pseudo-time for genes encoding components of the FGF signaling pathway (Fgf5, Fgf8, Fgfr1, Fgfr2, Spry4) and the endoderm marker transcription factors Gata4, Sox7 and Sox17 during EPI and PrE lineage specification. **e,** Gene expression patterns of FGF signaling pathway components, Gata and Sox transcription factor genes. Orange, black and green arrowheads point to high expression in ParE, VE and EPI, respectively. **f,** Dynamics of TF ratios as lineages are specified: Gata6/Nanog and Gata6/Fgf4 along EPI; Nanog/Gata6 and Fgf4/Gata6 along PrE, compared to changes in differential potential (dotted line). TF ratios were computed for each cell by using the MAGIC imputed data for each gene.

**Extended Data Figure 6:**
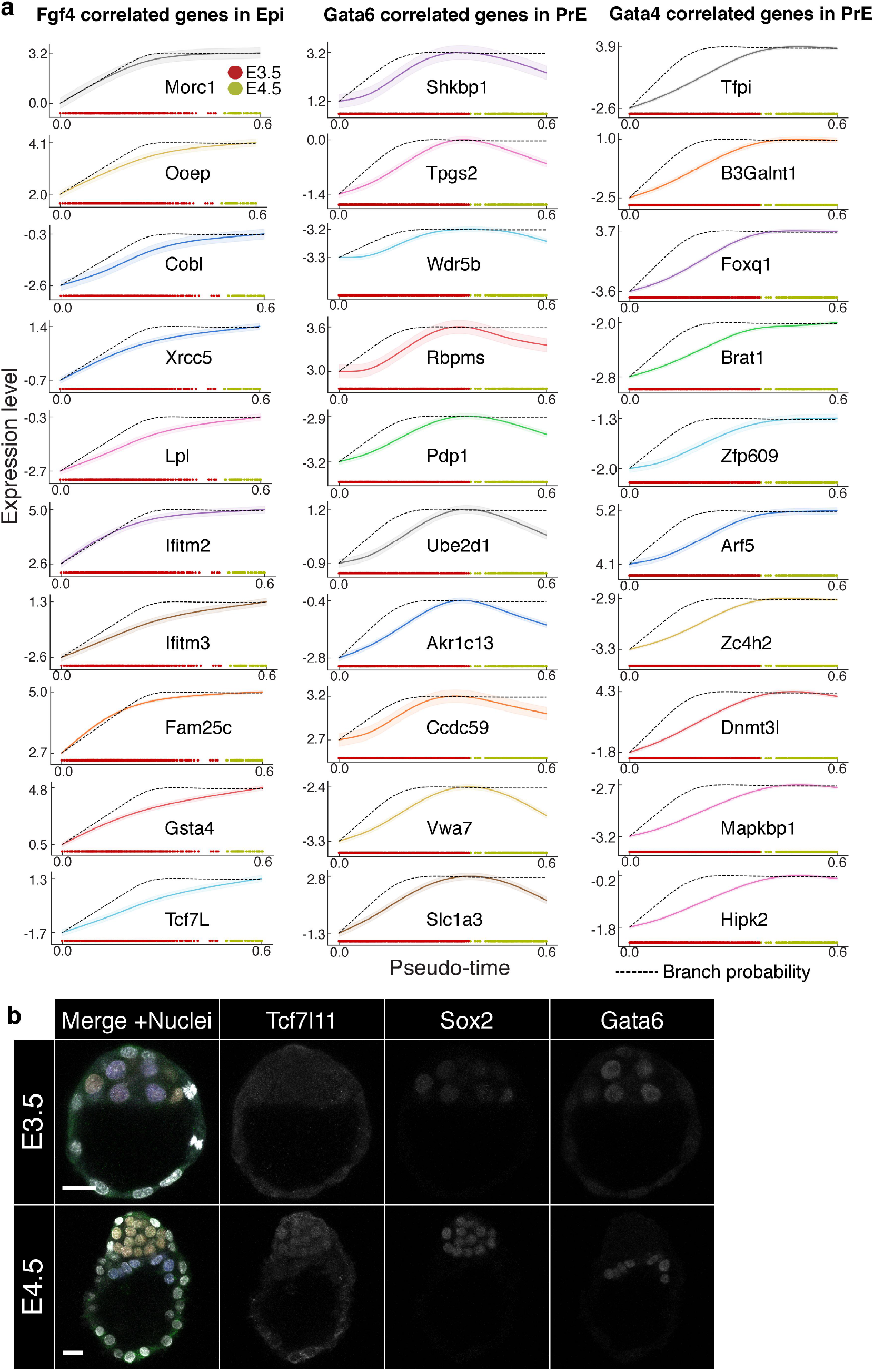
Expression trends of genes correlated with key lineage transcription factors. **a,** Gene expression trends for genes correlated with Fgf4 in EPI, and Gata4, Gata6 in the PrE. All correlations are > 0.97. **b,** Laser confocal images of Tcf7l1 protein expression at E3.5 (top panel) and E4.5 (bottom panel) Sox2 and Gata6 were used as EPI and PrE cell lineage markers, respectively.

**Extended Data Figure 7:**
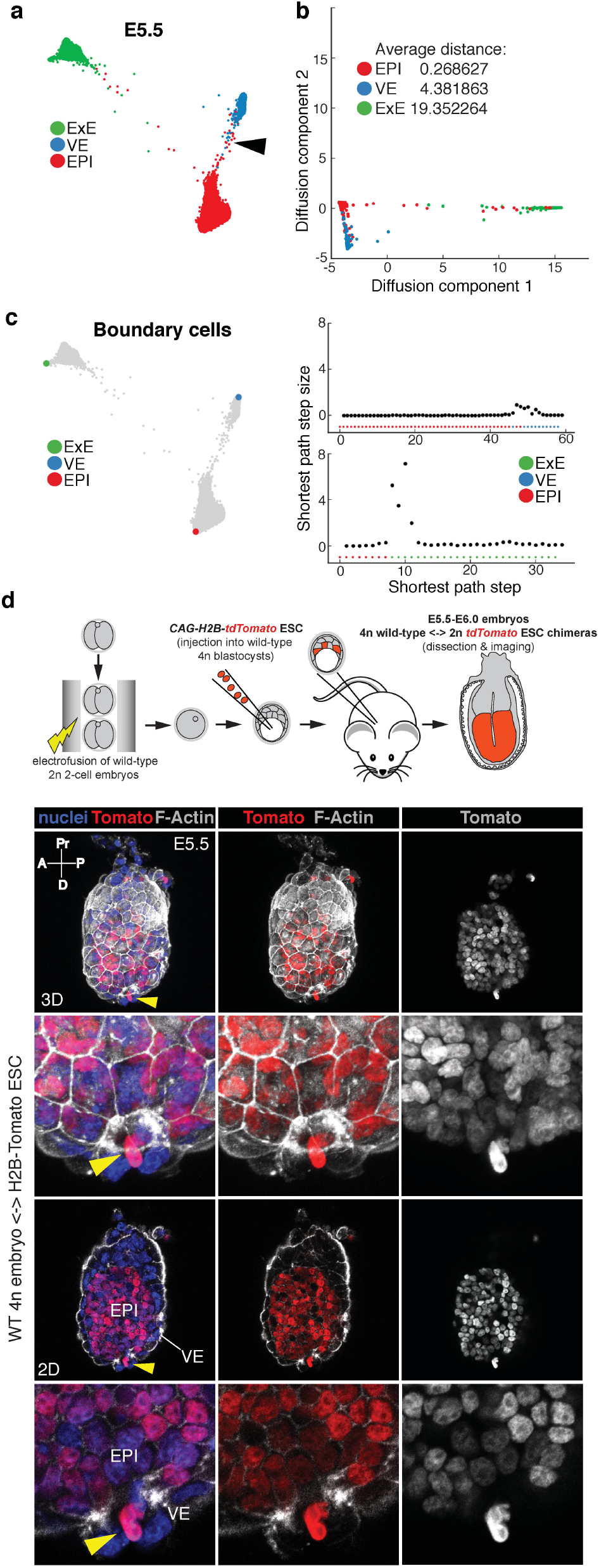
Force directed layouts of single E5.5 cells reveal relationships between EPI, VE and extra-embryonic ectoderm (ExE) lineages. **a,** Force directed layouts of E5.5 data generated after pooling replicates, showing the relationship between EPI, VE and extra-embryonic ectoderm (ExE) lineages. Cells are colored by cell type. Black arrowheads mark cells that transdifferentiate from EPI to VE. **b,** Plot showing the projection of EPI, VE and ExE cells along the first two diffusion components. **c,** Left panel: Plots highlighting the boundaries of the phenotypic space for each lineage identified as extremes of the diffusion components. Right panel: Plots showing the shortest path step sizes for paths from EPI to VE (top) and EPI to ExE (bottom). **d,** Laser confocal images of an E5.5 wild-type 4n <-> H2B-tdTomato embryonic stem cell (ESC) embryo chimera. An epiblast (EPI) cell is intercalating into the visceral endoderm layer (VE) (yellow arrowheads). Top two rows: Low and high magnification (3D images, maximum intensity projections) of an E5.5 wild-type 4n <-> H2B-tdTomato ESC embryo chimera. Bottom rows: Low and high magnification views (2D images) of an E5.5 wild-type 4n <-> H2B-tdTomato ESC embryo chimera. Embryo is counterstained with Hoechst to label nuclei, and Phalloidin to label F-Actin. A, anterior; D, distal; P, posterior; Pr, proximal.

**Extended Data Figure 8:**
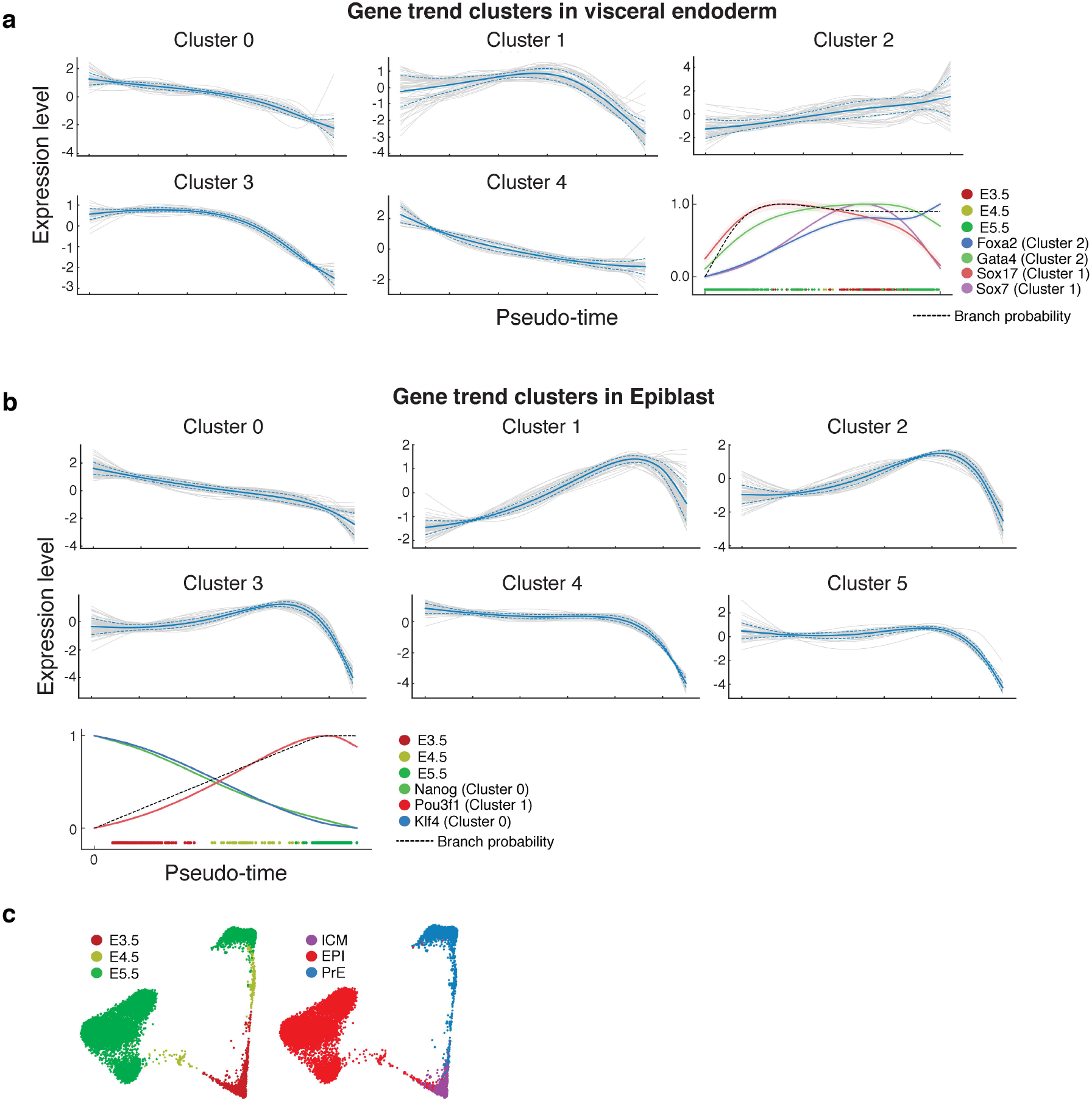
Differentiation of E3.5 ICM cells to PrE/VE and EPI lineages. **a,** Clustering of gene expression trends along visceral endoderm pseudo-time from E3.5-E5.5. Each grey line represents the gene expression trend of a particular gene. Solid blue line shows the mean expression trend of genes in the cluster, and dotted lines represent the standard deviation. Bottom right panel shows the gene expression trends of endoderm markers Foxa2, Gata4, Sox17 and Sox7 along pseudo-time from E3.5-E5.5 for the PrE/VE lineage. **b,** Clustering of gene expression trends along epiblast lineage pseudo-time from E3.5-E5.5. Bottom left panel shows the gene expression trends of epiblast markers Nanog, Pou3f1 and Klf4 along pseudotime from E3.5-E5.5. **c,** Force directed layouts of E3.5-E5.5 data showing the relationships between the PrE/VE and EPI lineages, excluding cells that trans-differentiate from EPI to VE.

**Extended Data Figure 9:**
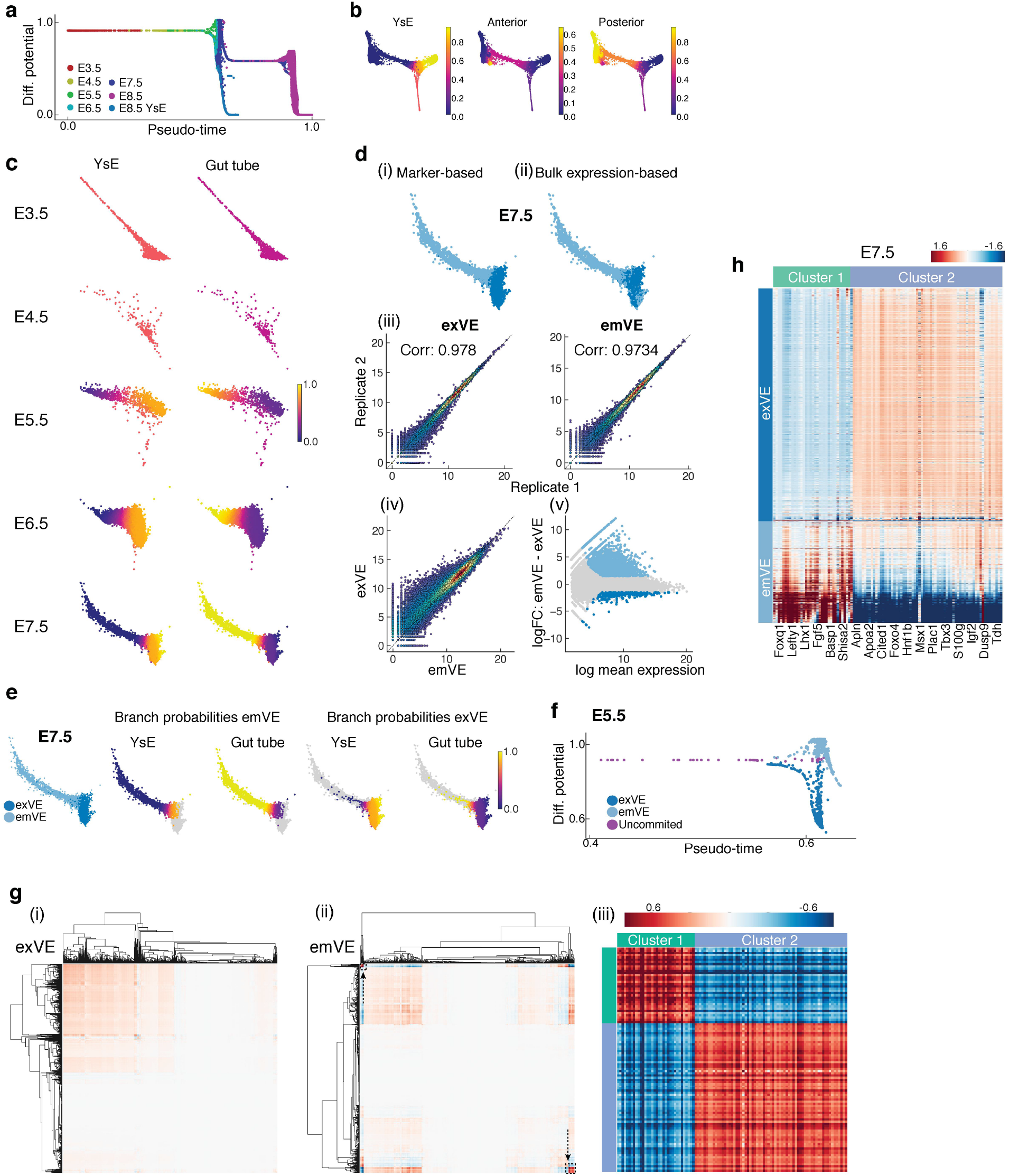
Emergence of spatial patterning of the embryo at E5.5. **a,** Plot showing Palantir pseudo-time versus differentiation potential of VE cells from stages E3.5-E8.75. Drops in differential potential occur at two time points. The first at E5.5, as cells acquire a distal versus proximal fate and the second at E7.5 as cells acquire an anterior versus posterior fate. **b,** Plots of branch probabilities of commitment towards yolk sac endoderm (YsE), anterior and posterior gut endoderm. **c,** Plots show the branch probabilities separately for cells at each time point. Anterior and posterior probabilities were summed to derived gut tube probabilities. **d,** Marker based (i) and bulk RNA-seq based (ii) prediction of exVE and emVE at E7.5. (iii) Plots showing the correlation between bulk RNA-seq replicates of exVE and emVE. (iv-v) Plots showing differentially expressed genes between of exVE and emVE derived using bulk RNA-seq data. **e,** Plots showing the branch probabilities of E7.5 exVE and emVE cells to commit towards YsE and gut tube. **f,** Plot showing pseudo-time versus differentiation potential of endoderm cells at E5.5 colored by the inferred cell type. (A zoomed in view of Extended Data Fig 9a). **g,** Gene expression covariance matrices of E5.5 exVE and emVE (i-ii). Cells are ordered by pseudo-time within each timepoint. Covariances were computed using MAGIC imputed data. A subset of genes that strongly covary with each other but are anti-correlated to genes in the other cluster are highlighted (black rectangles/arrows). (iii): Heatmap of the covariance of the highlighted gene clusters at E5.5 (see Supp. Table. 4). **h,** Heatmaps of highly expressed genes specifically in exVE or emVE at E5.5 also distinguish exVE and emVE cells at E7.5.

**Extended Data Figure 10:**
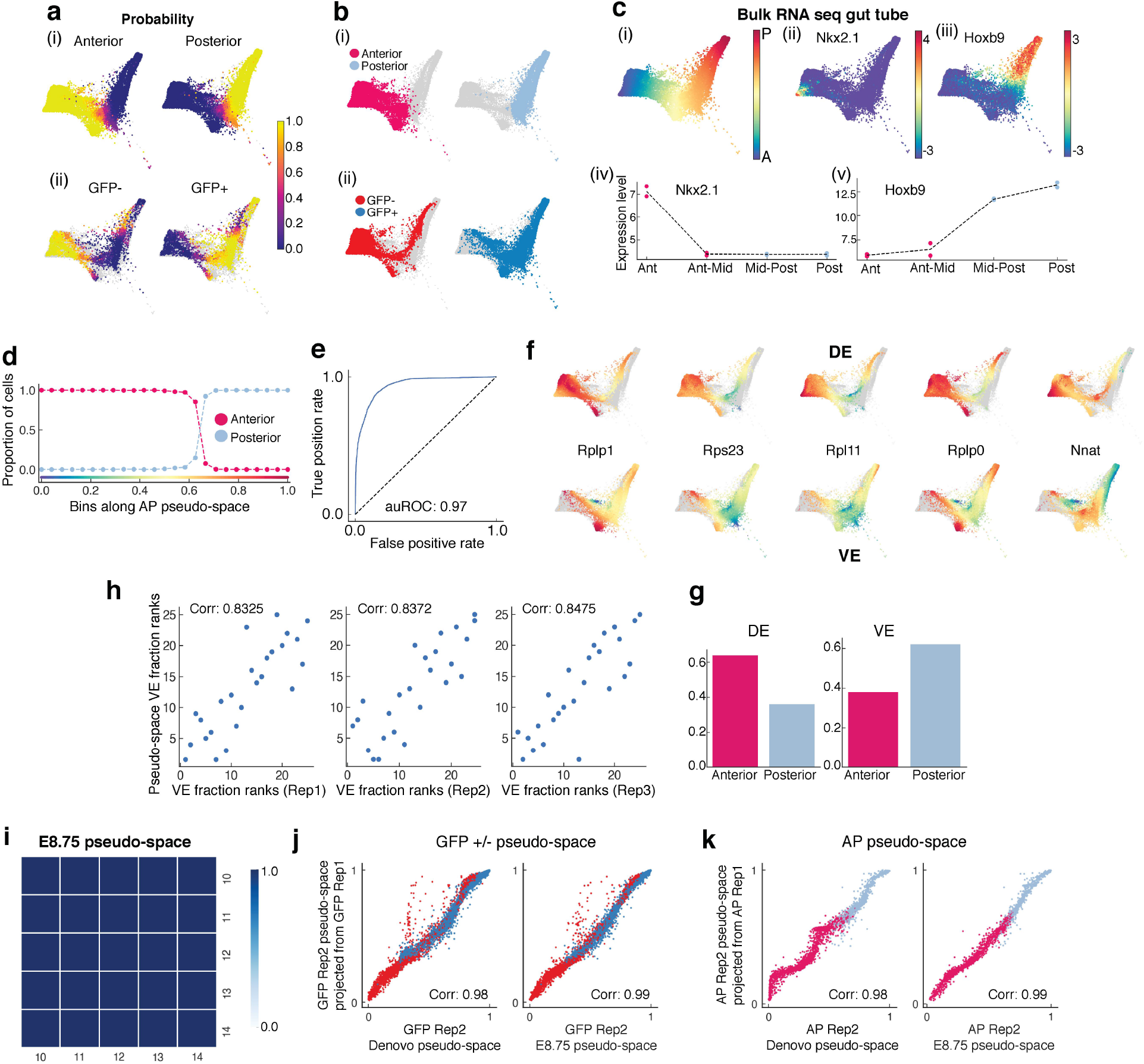
Characterization of E8.75 gut tube anterior-posterior pseudospace. **a,** Force-directed layout as in **Fig 6.** (i): Plots showing the probabilities of anterior-posterior positioning for the Afp-GFP-positive/Afp-GFP-negative cells inferred using the manifold classifier trained on anterior-posterior cell. (ii): Plots showing the probabilities of GFP-positive/GFP-negative status for the cells from the anterior-posterior compartment inferred using the manifold classifier trained on GFP-positive/GFP-negative cells. **b,** (i): Anterior and posterior positions of Afp-GFP-positive/AFP-GFP-negative cells inferred using probabilities in (a-i). (ii): GFP-positive/GFP-negative status of the anterior-posterior compartment cells inferred using probabilities in (a-ii). **c,** (i) : Plot showing the first diffusion component of the E8.75 cells. (ii-iii): Plots showing the expression of anterior marker Nkx2.1 and posterior marker Hoxb9 in E8.75 cells. (iv-v): Bulk RNA-seq expression of Nkx2-1 and Hoxb9 in quadrants of the gut tube along the AP axis compares with A-P single cell expression patterns. **d,** Plot showing the proportion of inferred and measured anterior and posterior cells in bins along the AP pseudo-space axis. **e,** Receiver operating curve for classification of E7.5 VE and DE cells. **f,** Plots showing the expression patterns of genes that are best predictive of the DE class in the VE-DE classifier (top - DE; bottom -VE). **g,** Bar plot showing branch probabilities of anterior and posterior positioning at the start of the pseudo-time ordering of DE cells (left) and VE cells (right) from E7.5 to E8.75. **h,** Plots comparing the ranks of proportion of GFP-positive cells along AP positioning in the Afp-GFP embryo-derived Neurolucida gut tube replicates (x-axis), and the ranks of VE cell proportions in bins along the AP pseudo-space axis (y-axis). **i,** Heatmap showing correlations between AP pseudo-space orderings determined using a varying number of diffusion components highlighting the robustness of the ordering. **j,** Plots comparing the AP pseudospace ordering of GFP-positive/GFP-negative cells (replicate 2) generated *de novo* using only the replicate 2 cells (x-axis, left) with the projected ordering from replicate 1 (y-axis). Right panel shows a similar comparison with the pseudo-space ordering determined using cells of both the replicates on the x-axis. **k,** Same as **j,** for replicates of anterior-posterior cells.

**Extended Data Figure 11:**
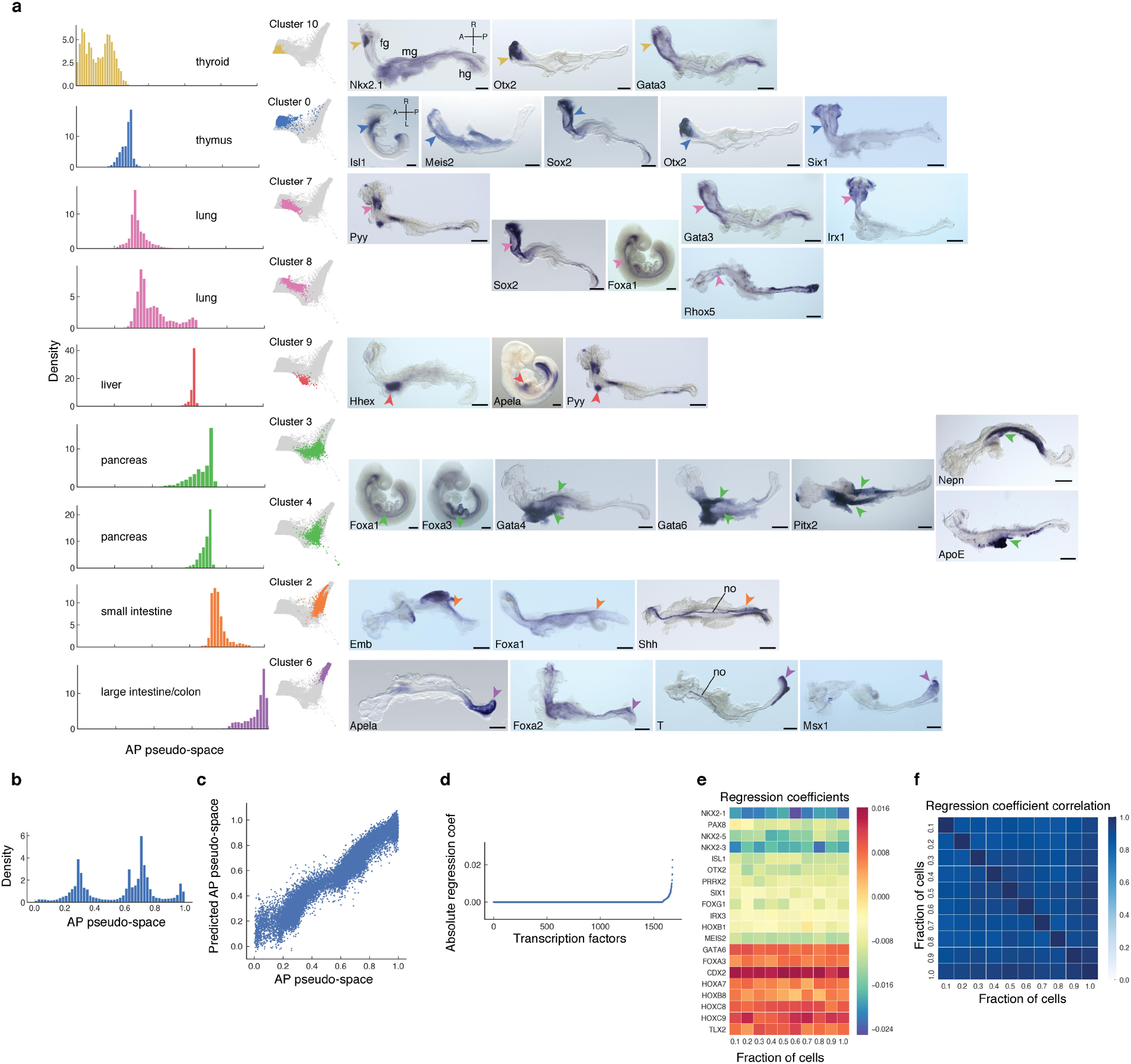
Spatial patterning of the gut tube at E8.75. **a,** Plots showing individual Phenograph clusters densities of the E8.75 gut tube cells ordered along AP pseudo-space (left panel) and in force directed layouts (middle panel). Whole-mount mRNA in situ hybridization of representative differentially expressed genes in each cluster on whole E8.75 embryos or micro-dissected E8.75 gut tubes (right panels). Arrowheads point to expression of representative gene for each particular cluster. All scale bars: 200μm, except for Nkx2-1: 100μm. A, anterior; fg, foregut; hg, hindgut; L, left; mg, midgut; no, notochord; R, right; P, posterior. **b,** Density of E8.75 cells along the AP pseudo-space axis. **c,** Comparison of empirical AP pseudo-space axis and the predicted AP pseudo-space using expression of TFs. d, Plot showing the ranking of different TFs according to their predictive power based on the regression model. **e,** Heatmap showing the coefficients for the top TFs when different proportions for cells are subsampled for the regression. **f,** Heatmap showing the correlation of TF coefficients in (e), highlighting the robustness of TF coefficients in regression.

**Extended Data Figure 12:**
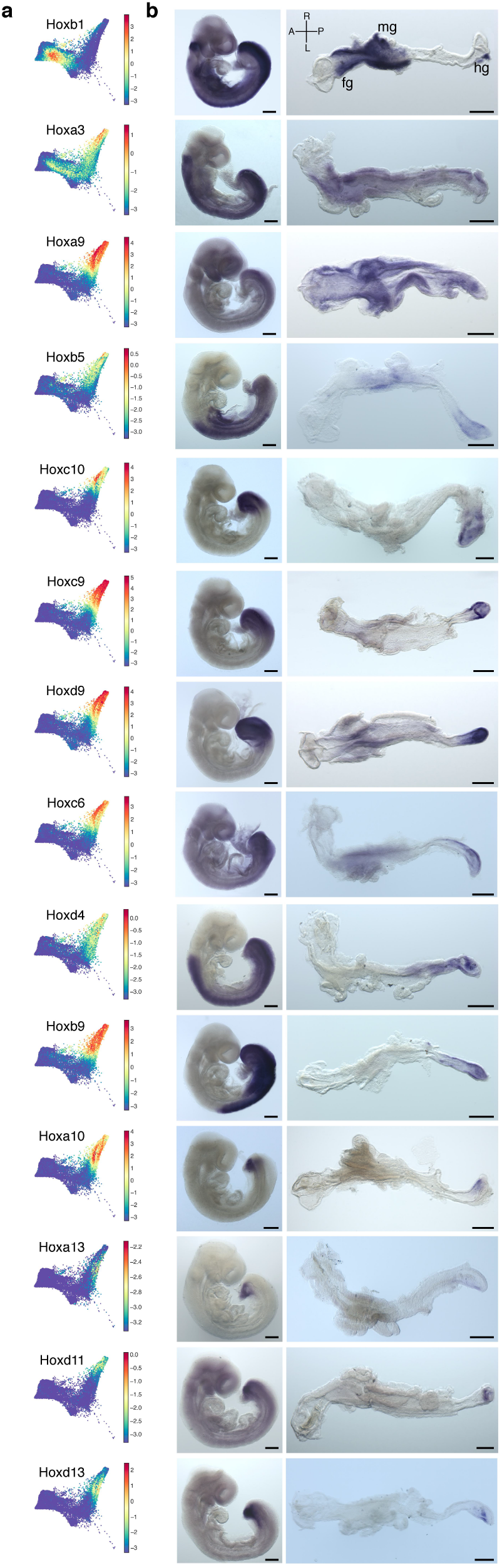
Hox gene expression within the E8.75 gut tube. **a,** Force directed plots of the expression of Hox genes expressed by gut endoderm cells at E8.75. **b,** Whole-mount mRNA in situ hybridizations on whole E8.75 embryos and microdissected gut tubes of Hox genes depicting their distribution along the AP axis. All scale bars: 200μm, except for Hoxc10, Hoxd11: 100μm. A, anterior; fg, foregut; hg, hindgut; L, left; mg, midgut; R, right; P, posterior.

## Supplementary Figures

**Supplementary Figure 1:**
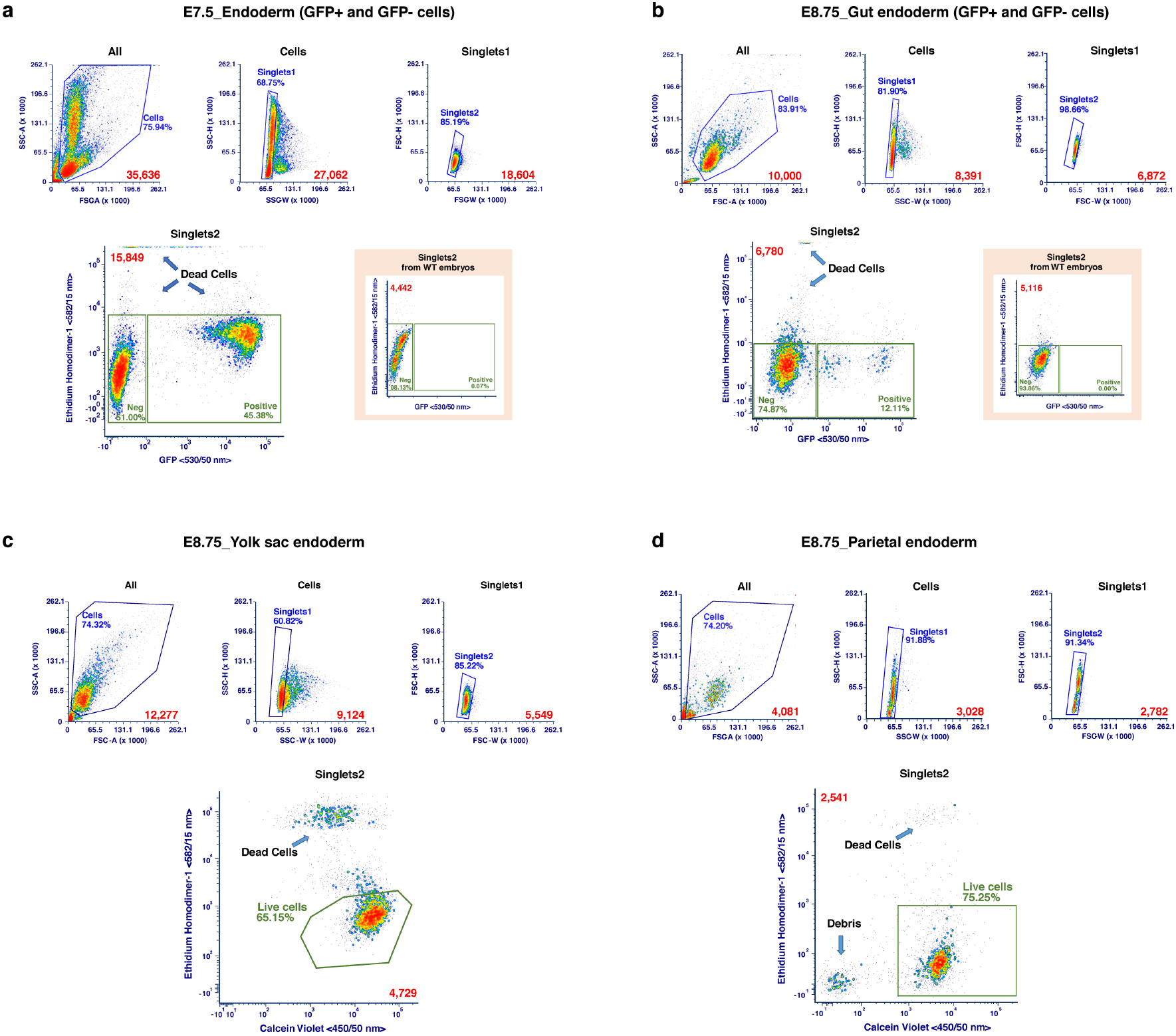
Gating strategy for sorting endoderm cells from mouse embryos. **a,** Gating strategy used to sort single E7.5 endoderm cells. Top: Aggregates were excluded with a SSC (Height vs. Width, Singlets1) followed by a FSC (Height vs. Width, Singlets2) plot. Bottom left plot: The final singlets population (Singlets2) was plotted on an Ethidium Homodimer-1 (582/15 nm) vs. GFP (530/50 nm) to exclude dying cells and sort GFP-negative and GFP-positive cells. Bottom right plot: GFP negative control. Numbers in red are the total numbers represented in each plot. **b,** Gating strategy used to sort single E8.75 gut tube cells. Top: Aggregates were excluded with a SSC (Height vs. Width, Singlets1) followed by a FSC (Height vs. Width, Singlets2) plot. Bottom left plot: The final singlets population (Singlets2) was plotted on an Ethidium Homodimer-1 (582/15 nm) vs. GFP (530/50 nm) to exclude dying cells and sort GFP-negative and GFP-positive cells. Bottom right plot: GFP-negative control. Numbers in red are the total numbers represented in each plot. **c,** Gating strategy used to sort single E8.75 yolk sac cells. Top: Aggregates were excluded with a SSC (Height vs Width, Singlets1) followed by a FSC (Height vs Width, Singlets2) plot. Bottom: The final singlets population (Singlets2) was plotted on an Ethidium Homodimer-1 (582/15 nm) vs Calcein Violet (450/50 nm) to exclude dying cells and debris and sort only viable live cells. Numbers in red are the total numbers represented in each plot. **d,** Gating strategy used to sort single E8.75 parietal endoderm cells. Top: Aggregates were excluded with a SSC (Height vs. Width, Singlets1) followed by a FSC (Height vs. Width, Singlets2) plot. Bottom: The final singlets population (Singlets2) was plotted on an Ethidium Homodimer-1 (582/15 nm) vs. Calcein Violet (450/50 nm) to exclude dying cells and debris and sort only viable live cells. Numbers in red are the total numbers represented in each plot.

**Supplementary Figure 2:**
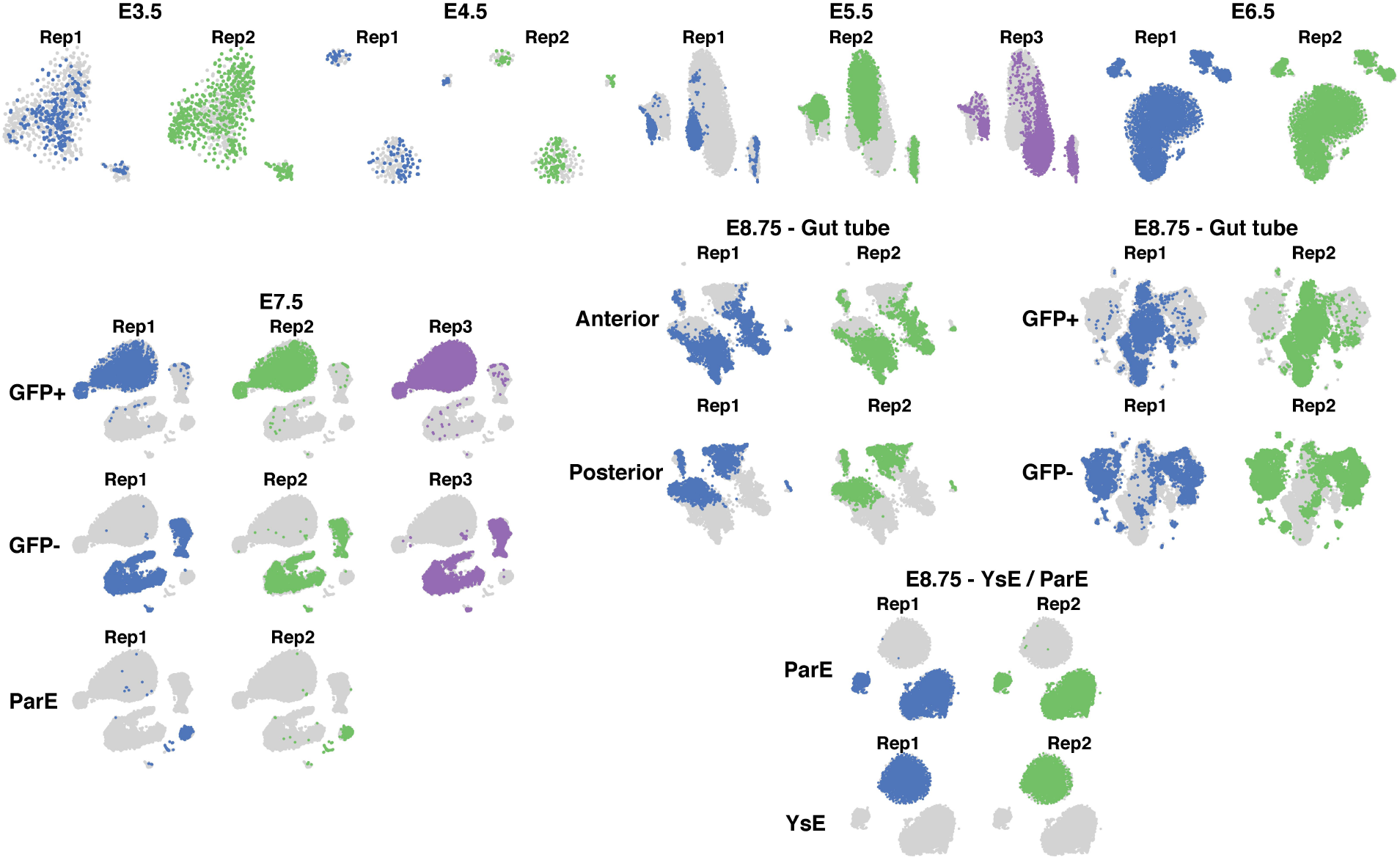
tSNE plots of all single cell libraries generated and stages of mouse embryo. tSNE plots for each tissue and embryonic stage, cells color-colored by replicate. The following cells were collected from the different stages: E3.5-E5.5: Whole embryos, E6.5: Visceral endoderm cells, E7.5: GFP-positive visceral endoderm, GFP-negative definitive endoderm and parietal endoderm cells, E8.75: cells from anterior and posterior halves of gut tube, GFP-positive visceral endoderm, GFP-negative definitive endoderm, yolk sac (YsE) and parietal endoderm (ParE) cells. Replicates are color-coded.

